# High Resolution Nanostructure with Two-stages of Exponential Energy Dissipation at the Ultrathin Osteochondral Interface Tissue of Human Knee Joint

**DOI:** 10.1101/2021.10.18.464899

**Authors:** Xiaozhao Wang, Junxin Lin, Zonghao Li, Yuanzhu Ma, Xianzhu Zhang, Qiulin He, Qin Wu, Wei Wei, Xudong Yao, Chenglin Li, Wenyue Li, Shaofang Xie, Yejun Hu, Shufang Zhang, Yi Hong, Xu Li, Weiqiu Chen, Wangping Duan, Hongwei Ouyang

## Abstract

Cartilage adheres to subchondral bone *via* a specific osteochondral interface tissue where forces are transferred from soft cartilage to hard bone without fatigue damage over a lifetime of load cycles. However, the fine structure and mechanical properties of osteochondral interface tissue remain unclear. Here, we identified an ultrathin ∼20-30 μm calcified region with two-layered micro-nano structures of osteochondral interface tissue in human knee joint, which exhibited characteristic biomolecular compositions and complex nanocrystals assembly. Within this region, an exponential increase of modulus (3 orders of magnitude) was conducive to the force transmission which was verified by finite element simulations. The nanoscale heterogeneity of hydroxyapatite, along with enrichment of elastic-responsive protein-titin which is usually present in muscle, endowed the osteochondral tissue with excellent energy dissipation and fatigue resistance properties. Our results provide potential design for high-performance interface materials for osteochondral interface regeneration and functional coatings.

## INTRODUCTION

Interface between soft and hard materials is often the weakest point. However, connections of soft-to-hard tissues in human body with an ultrathin transitional interface possess superior energy dissipation and exceptional fatigue-resistant capabilities, which are extremely difficult to replicate with a synthetic material.^1, 2^ Of these, osteochondral interface in human knee joint, allowing robust attachment of compositionally, structurally and mechanically mismatched materials-articular cartilage and subchondral bone, and enabling effective heavy force transfer between soft and hard tissues over a lifetime of load cycles, is the most sophisticate interface model for learning.^3^

Osteochondral interface resists exceedingly high stresses under harsh mechanical environment. The moduli between cartilage and subchondral bone differed by thousand-fold,^4^ which makes osteochondral interface tissues subject to large stress concentrations and subsequently leads to tissue failure.^5^ However, osteochondral interface with load-bearing function shows durability, and physically works properly during the whole lifespan. Its secret of success should be their sophisticated structures with functionally graded designs whose properties far exceed a simple mixture of their isolated components.^6, 7^ The intermediate change of stiffness within the interface between articular cartilage and subchondral bone also make contributions.^8^ Osteoarthritis is commonly accompanied by the breakdown of osteochondral interface, and load transfer restoration of injured osteochondral interface poses a great challenge to date.^3, 9^ For this reason, identifying the structural and mechanical mechanism behind the natural minimization of stress concentration at osteochondral interface is important for understanding the pathogenesis and treatment of osteoarthritis.

The osteochondral interface is described as a graded calcified cartilage region with discontinuous unmatched collagen-I and collagen-II fibers, distributed from mineral-rich into overlying mineral-free cartilaginous tissue, where proteoglycan matrix and carbonated hydroxyapatite similar to bone is detected.^10, 11^ Generally, spatial gradients of components comprising minerals and organic molecules with a particular direction, are beneficial for load transfer through their extrinsic toughening mechanism of effective energy dissipation.^7, 12, 13^ However, the mixture and assembly of compositions referred to the current findings could not precisely explain the superior mechanical property of osteochondral interface tissues.^10, 11, 14^ The high-resolution analysis based on the composition and structure of osteochondral interface tissue could provide paradigm and rational design for future development of high-performance soft-hard composite materials.

Here, by using materials science approaches, we investigated the two-layered micro-nano structure variations, biomolecular compositions and mineral assembly of the osteochondral interface of human knee joint at multiscale. The detailed stiffness transition of the interface tissue was evaluated by a nanoindentation method, and further simulated with finite element analysis (FEA) to validate its superiority for force transfer. The underlying mechanism on how the intricate interplay of compositions and structures determined the exceptional mechanical responses of osteochondral interface tissues was proposed in the end.

## RESULTS AND DISCUSSION

### An Ultrathin Osteochondral Interface with Two-layered Microstructures

We first examined the microstructure of human cartilage tissue with synchrotron radiation microtomography (SR-μCT), scanning electron microscopy (SEM) and scanning transmission electron microscopy (STEM). Schematic illustration highlighted the complete architecture of the cross-sectioned cartilage tissue including articular cartilage (AC), osteochondral interface tissue and subchondral bone (SB) (Figure 1a, Data Figure S1). The osteochondral interface tissue between AC and SB was visualized with the distinct tidemark pattern as the frontier (Figure 1b, c), matched well with the histological staining results and atom force microscopy topographies (Figure Data S2), revealing the onset of the calcified interface.^15^ The density dependent colour electron micrographs (DDC-SEM) of osteochondral interface tissues present graded mineral distributions with distinct morphologies (Figure 1d, details see Data Figure S3). Corresponding elemental analysis by SEM with energy-dispersive X-ray spectroscopy (EDX) mapping showed the distributions of calcium (Ca) and phosphorus (P) elements (Data Figure S4), illustrating smooth grading. Contents of Ca and P across the interface showed exponential increment profiles (Figure 1e), further highlighting the transitional zone from mineral-free to mineral-rich region. Therefore, this region was defined as osteochondral interface and the corresponding ultrathin layer was spanning over ∼20-30 μm for all samples (Figure 1f). The micrographs of the interface (enlarged images from the yellow box in Figure 1d) revealed the nature of the gradually mineralized CaP particles assembled in organic matrix. Three dominant mineral structures were present as the combinations of small dense mineral granules, more and larger needle-like globules and mature platelets composed of small spherical particles (Figure 1d, g, h), and the latter is similar to the apatite crystallites of natural bone.^16^ The spherical particles are considered to be poorly matured mineral structure and may serve as the transient precursor for further mineral maturation.^17, 18^ With the emergence of nanoparticles and mineral maturation at the interface, STEM images further demonstrated that porous structure of 100-400 nm diameter occupying spaces between collagen fibrils and minerals, gradually became smaller, and finally vanished to be densely packed minerals (Figure 1h). Thus, two-layered micro-nano structure variations, between a porous layer near AC side and dense layer near SB side, were exhibited at the osteochondral interface. The porous layer was suggested to facilitate transportations of mineral ions and precursors to form isolated mineral particles,^19^ which is beneficial for smooth transition. The densely packed structures will block solute diffusion, thus preventing the vascular invasion from SB into AC and helping pressurization of AC to resist load.^20, 21^ This characteristic microstructure due to distinct mineral distributions at osteochondral interface contributed a lot to tissue functions. When we removed the mineral particles from the interface tissue, the original microstructures were altered, consequently inducing a gap at the interface (Data Figure S5). It may further imply the pivotal role of minerals distribution at the transitional interface in maintaining tissue integrity and helping compliant attachment of different tissues.^3, 22^ Similar gradient mineral distributions were also found on other connective tissues (tendons/ligaments to bone),^1, 23, 24^ which encouraged many biomimetic approaches to synthesize nanoparticle glues.^25–27^ Since spatial gradient in microstructure along the longitudinal direction between two dissimilar materials is suggested to alleviate stress concentration in joint move,^6, 7, 28^ it prompts us to further illustrate the detailed mechanical response of interface tissue.

**Figure 1.**
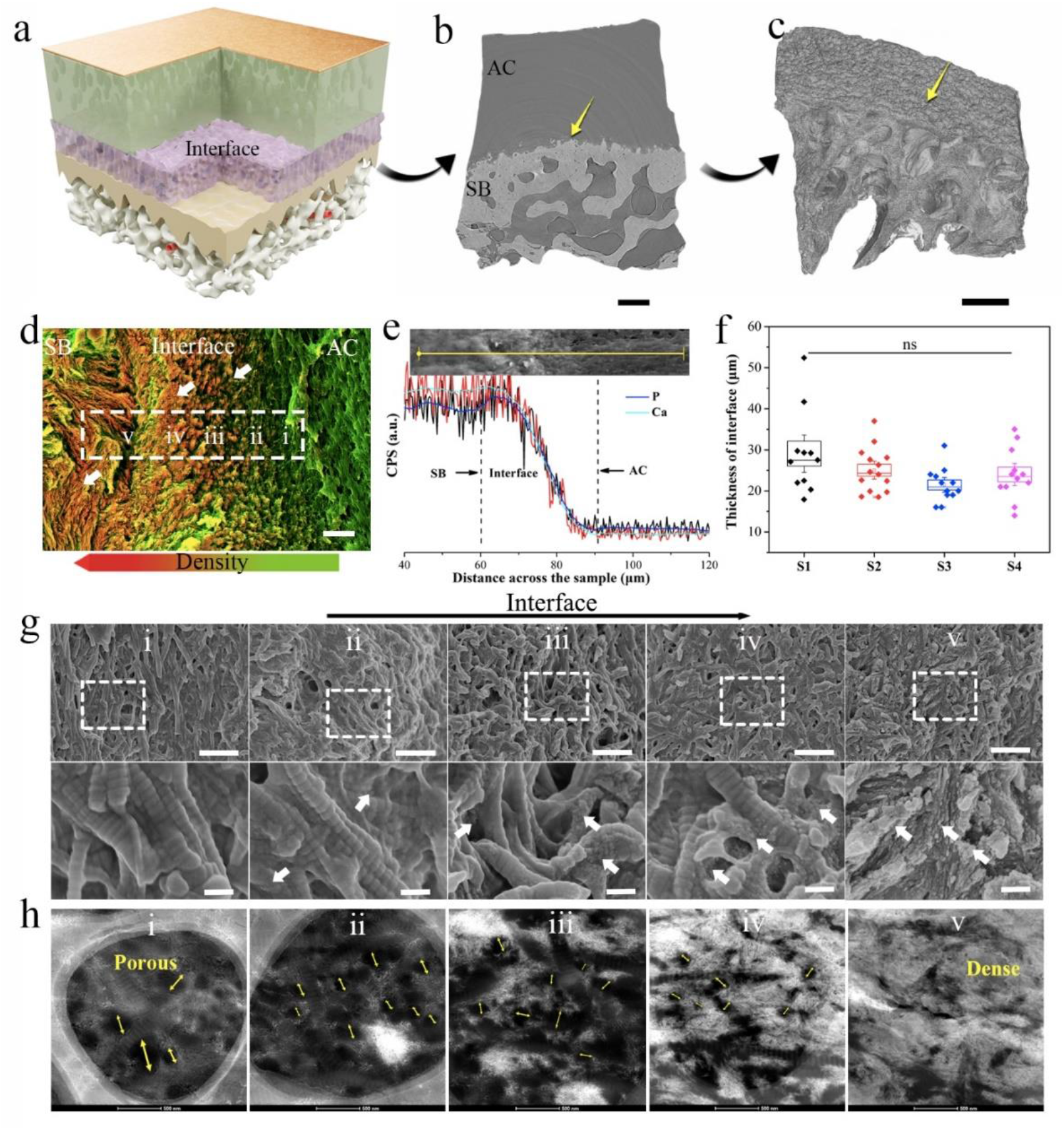
The microstructures variation of the osteochondral interface. **(a)**. Schematic illustration of human articular tissue showing osteochondral interface. **(b)**. Representative phase-contrast SR-μCT images of sectioned cartilage tissues showing the cartilage, osteochondral interface and SB. **(c)**. The reconstructed 3D volumetric representation of the structures including osteochondral interface and SB tissues. The yellow arrows in b and c highlighted the front of the mineralized interface tissues. **(d)**. DDC-SEM micrographs of osteochondral interface tissues presenting gradual mineral distributions with distinct morphologies, micrographs were colored in post-processing by combining images acquired by backscatter electron and secondary detectors (red-minerals, green-organic materials, the original images and processing details see Figure S3). White arrows indicated evolution of mineral morphologies at interface. (**e)**. SEM images and corresponding EDX line scan (Ca and P) (down) across the interface revealing the depth-dependent contents of Ca and P. **(f)**. The thickness of the calcified interface of human knee joint samples quantified by EDX line scan. **(g)**. SEM images showing the microstructures of the interface from the uncalcified region to maturely mineralized area (i-v in d). White arrows denoted the mineralized nanoparticles on collagen fibrils, indicating the mineralization was gradually mature throughout the interface. **(h)**. STEM images of transverse sections of osteochondral tissue at different site presenting the transition of porous structure (indicated with yellow arrows) to dense structure and particles distributions throughout the interface. Scale bar in b-c: 50 μm, d: 5 μm, g: 1 μm and 200 nm (down), h: 500 nm.

### Exceptional Modulus Changes within the Ultrathin Osteochondral Interface Tissue

To understand the mechanical response of osteochondral interface, the nanoindentation test was performed on the transverse section of cartilage samples comprising AC, osteochondral interface and SB region (Figure 2a-d). Spherical indenter heads were used in the test to avoid tissue damage and Hertz model was applied to calculate the elastic modulus of different tissue sites.^4^ Tissue modulus differs greatly (three orders of magnitude, varied from 3.9 MPa to 1.87 GPa) over the micrometer-thick interface, consistent with previous studies.^8, 22^ An obvious wavy band of lower modulus parallel to tidemark border near AC region was identified as the front of the interface. We divided tissue modulus increments into two stages. In stage 1 (near AC region), an order increment of magnitude of modulus was revealed over the first ∼5-10 μm of transitional region, and two orders of magnitude larger in modulus were demonstrated in stage 2 across the length scale of ∼10-20 μm (near SB region). The two-stages increment of tissue modulus was correlated well with the two-layered micro-nano structure variations (Figure 1g-h). The gradation in porosity and mineralization serves critical roles in driving tissue stiffness across the interface region.^7, 29^ In addition, the intermediate stiffness of interface tissue is beneficial for smooth transition between AC to SB.^22^

**Figure 2.**
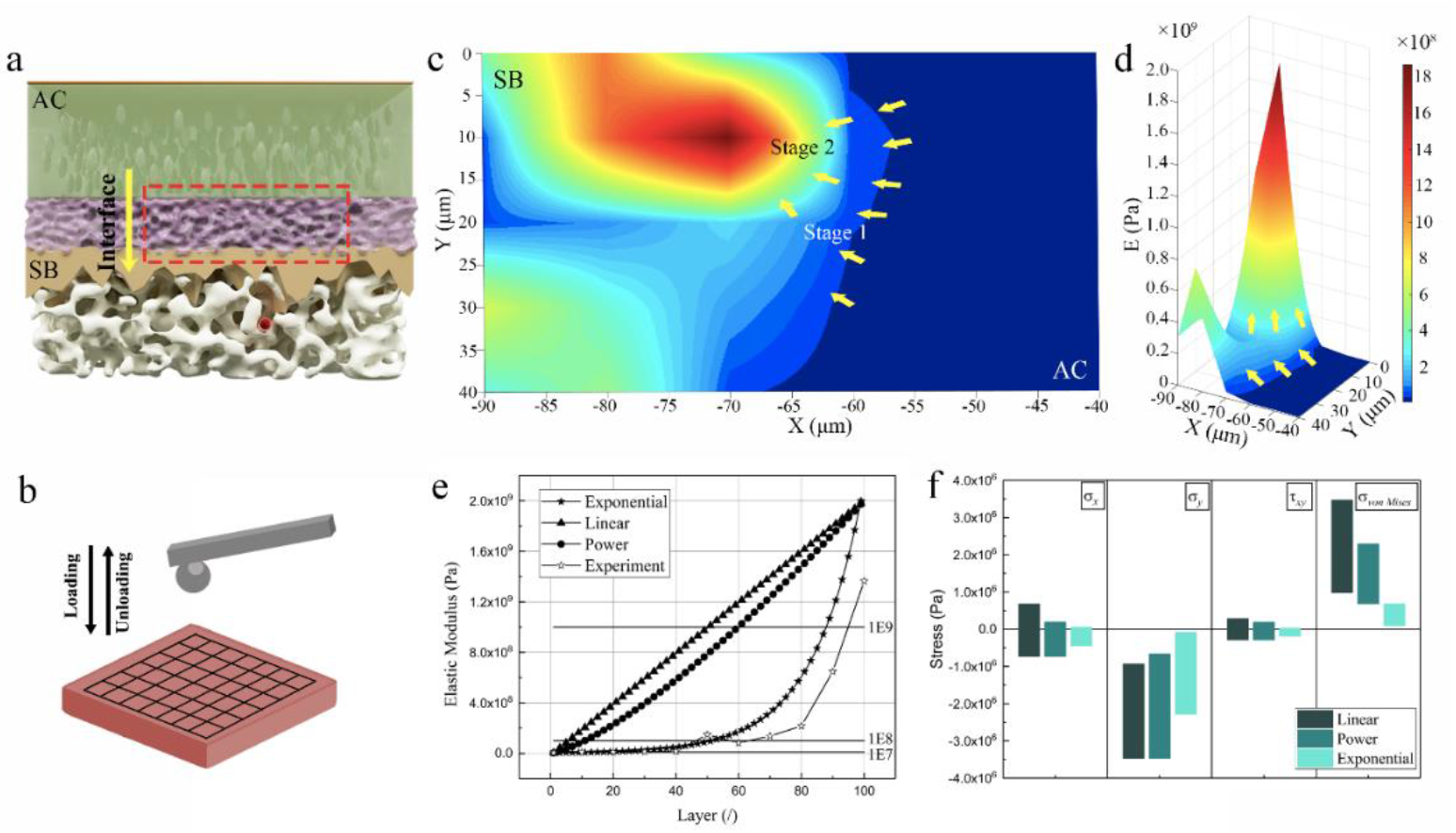
Biomechanics of the osteochondral interface. **(a)**. Schematic view of osteochondral interface, the red box was the region of tissue to be detected. **(b).** Schematic showing nanoindentation test of osteochondral interface. **(c-d).** Efficient modulus maps of the rectangular area in panel with a 10 μm spacing. Yellow arrows indicated two stages of modulus increments throughout the interface. The transition in mechanical increment coincided with the transition in local microstructure variation region. **(e).** Interlaminar modulus of elasticity. **(f).** The stress components and Mises stress of the interface tissue for different function types.

To further validate the mechanical superiority in gradient stiffness of the interface tissue, FEA was performed to simulate the mechanical behaviors of the interface tissue. Three mechanical models of elastic modulus at the interface were proposed, and the exponential function was examined to be the most fitting model corresponding to the experimental data (Figure 2e). To prove that such gradient modulus distributions of interface tissue could promote force transfer, the von-Mises stress distribution maps of the intact model were calculated (Data Figure S6). The results revealed that exponentially distributed modulus of interface tissue showed a much smaller high-stress region than the other function forms. The high-stress region was also observed to be closer to SB tissue with higher modulus, while the obviously decreased stress emerged on AC side with lower modulus. It should be resulted from the fact as: stage 1 is soft and prone to deform for energy dissipation and relieve stress concentration, and stage 2 is hard enough to resist against crack passing; together they lead to efficient force transfer without damage.^30–33^ In addition, the maximum stress in the interface tissue was significantly reduced (Figure 2f), which was largely ascribed to the graded mechanical properties of tissue.

These results highlight that exponential increase in tissue modulus of the interface tissue is ideal for load transfer from soft AC tissue to hard SB tissue, which avoid tissue failure caused by high stress on AC side. A recent study has reported that high-load induced fracture is rarely formed within healthy osteochondral interface tissue, while that is common in diseased osteochondral interface in osteoarthritis joint, highlighting normal osteochondral interface is superior in load transfer.^9, 34^ Such superiority largely depends on the two-layered micro-nano microstructure variation and exponential mechanical response of interface tissue. In addition, local arrangement of constituents is another nonnegligible material motif.^7, 35, 36^

### The Nanoscale Heterogeneity of HAp Assemblies across the Osteochondral Interface Tissue

Gradated transitions in composition across the ultrathin interface, especially the heterogeneity between the two dissimilar materials, could redistribute thermal stresses and thereby influence the mechanical properties of the interface.^35^ To deeply understand this, we next identified the high-resolution compositions and their spatial distributions at the microscale and nanoscale. Mineral crystals at the osteochondral interface have been reported to show similar size compared with that of bone,^10^ but their detailed crystalline compositions, mineral-to-matrix distribution as well as ordered assembly of mineral particles are still elusive. Here, we first quantitatively defined the mineral phase of CaP at the interface. Ca/P ratios at various sites across the interface ranged from 1.45 (interface site) to 1.76 (SB region) (Data Figure S7 a-d), which was similar to that of natural bone.^37^ This observation revealed that immature features of CaP at the interface which quite coincided with the granule and globule-like morphologies of nanoparticles.^16–18^ XRD results revealed that the mineral phase of the interface was mainly hydroxyapatite (HAp) with poor crystallinity (Data Figure S7e).^38^ Raman spectra of the interface exhibited the dominated peak of PO_4_^3−^ *v*1 symmetric stretch at 960 cm^−1^ and a weaker peak of CO_3_^2−^ *v*1 in plane vibrations at 1071 cm^−1^, indicating the mineral at the interface was carbonate-substituted HAp (Figure 3a).^39^ As the position and shape of PO_4_^3−^ and CO_3_^2−^ band at 960 cm^−1^ and 1071 cm^−1^ reveal mineral crystallinity, relative amounts of components and ionic substitution, we further produced compositional maps for HAp and matrix, and demonstrated their spatial distributions across the interface. An increase of relative HAp, substituted carbonate content as well as mineral-to-matrix ratio were apparently observed at the interface, indicating that carbonate-substituted HAp showed a depth-dependent distribution (Figure 3b-d). The crystallinity of HAp across the interface, shown by the full-widths at half-maximum (FWHM) of the peak at 960 cm^−1^, was gradually increased from AC to SB region (Figure 3e), while the distribution of CO_3_^2−^/PO_4_ ratio showed a converse trend (Figure 3f), highlighting the presence of abundant carbonate-substituted HAp with poor crystallinity at the frontier region of interface near AC where spherical minerals were mainly observed (stage 1). The region with higher crystallinity and lower carbonate substitution of HAp was found in stage 2, which exhibited obviously densely packed structures. In other words, as carbonate substitution ratios decreased across the interface region, increased crystallinity and microstructure change of minerals appeared and correlated well with the elevated tissue moduli. A recent study has shown that carbonate substitution is sufficiently to control the mineral morphology and mechanics of crystals.^40^ Increase of carbonate substitution ratios significantly reduces modulus of apatite and enamel. ^41, 42^ These results suggested a correlation between carbonate contents and mechanical properties of minerals at osteochondral interface. Thus, chemical composition (carbonate substitution) affects the mineral morphologies and maturity,^17^ subsequently dictating the mechanical properties.

**Figure 3.**
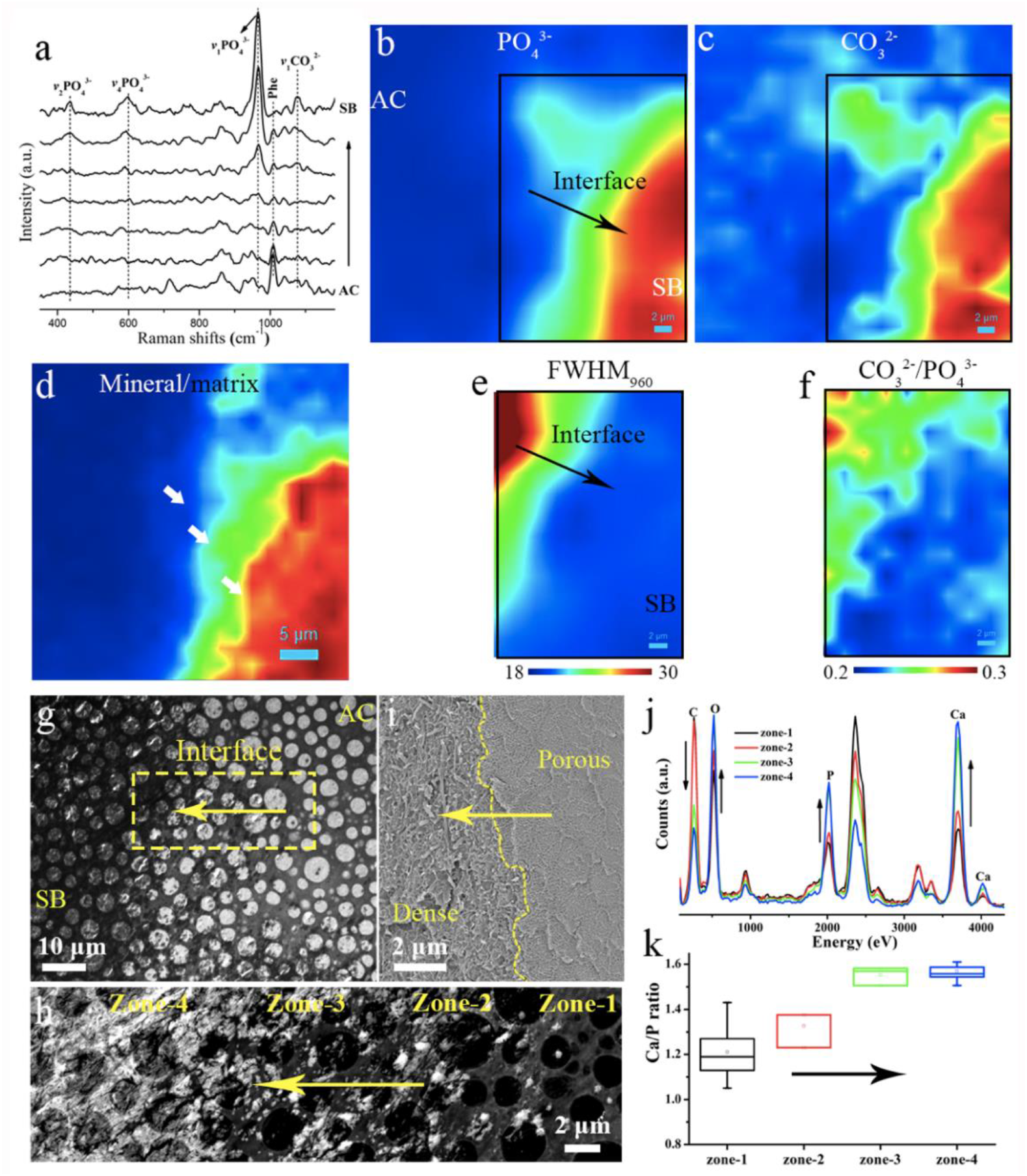
Compositional analyses of the osteochondral interface. **(a)**. Raman spectra of the interface from AC to SB. **(b-d)**. Raman peak intensity maps of PO_4_^3−^ *v*1 band at 960 cm^−1^, CO_3_^2−^ *v*1 band at 1071 cm^−1^ and mineral-to-collagen ratio (960 cm^−1^/1595-1700 cm^−1^), showing relative content and distribution of HAp. The gradient distribution of CO3^2−^ revealed the mineral phase of HAp over the interface was carbonate-substituted HAp. **(e)**. FWHM of the peak at 960 cm^−1^ presenting the mineral crystallinity across the interface. **(f)**. The maps of CO_3_^2−^/PO_4_ ratio revealing ionic substitution of minerals. The lower FWHM and CO_3_^2−^/PO_4_^3−^ ratio indicated that the HAp minerals were less substituted and more crystalline. **(g-h)**. TEM and High-angle annular dark-field (HAADF)-STEM images showing the morphologies of interface. **(i)**. Cryo-SEM images of the interface revealing the microstructure was changed from porous to dense. **(j)**. TEM-EDX spectra collected from different sites from zone 1-4 in h showing the elemental changes of the interface. **(k)**. Variation in Ca/P ratios of zone-1 to zone-4 calculated from EDX spectra. n = 9∼16 for each zone. Scale bar in a-b, d-e and i-h = 2 μm, in c= 5 μm, in g= 10 μm.

Transmission electron microscopy (TEM), cryo-SEM and STEM images further showed the spatial distributions of mineral nanoparticles over the interface with distinct morphologies (Figure 3g-i). If the interface is divided into four zones, a substantial increase of the content of Ca, P and O and a decrease of C content were observed over the interface region (Figure 3j). The Ca/P ratio within each zone spanned a wide range from 1.2 to 1.59 (Figure 3k, calculated by TEM-EDX), which was correlated well with the different degree of crystallinities observed in SEM-EDX and Raman results, further proving the progression of mineral maturation within the interface. Collectively, these results indicated that mineral particles underwent not only morphological variations, but also chemical composition changes at the interface region. Such microheterogeneity could be the potential cues for the excellent two stages mechanical performance of the interface.^24, 33^

To further elucidate the elaborate chemistry gradients between each nanoparticle with distinct morphologies, we also performed ultrastructural analyses for nanoparticles based on four zones throughout the interface marked i-iv. High resolution transmission electron microscopy (HRTEM) -STEM images confirmed the spherical, triangular and polygonal morphologies of the particles, with a dense core and needle-like periphery and growing size over the interface (Figure 4a). Higher Ca and P of different particles with lower C content were confirmed by EDX mapping analysis (Figure 4a), supporting the results in Figure 3j. Local chemical and atomic environment of different particles (i-iv) were further analyzed by electron energy loss spectroscopy (EELS). In each particle, the presence of C (k edge) and Ca (L_2,3_ edge) was confirmed, in varying contents; of note, similar content of C and Ca at the core and exterior region was identified in particle i; whereas, other particles (ii-iv) showed higher Ca and lower C content expanded from exterior to the core (Figure 4b). Bond assignments for each edge were included (Data Table. S1), in which no obvious changes in fine structures of Ca (L_2,3_ edge) were found between each particle, whereas that of C (k edge) showed striking difference. The carbon (k edge) spectra taken from all particles comprised four characteristic peaks: A at 284.5 eV, B at 287.2eV, C at 290.2 eV, and D at 297 eV (Figure 4b). The 1s-π* transition at around 284.5 eV (peak A) along with the transition peak at 297 eV (peak D) were attributed to the amorphous carbon, which was possibly ascribed to the embedding resin.^17^ The shoulder peak B originating from 1s-π* transition of the carbonyl group,^17, 43^ was predominantly observed in particle i-ii, and less intense in particle iii-iv. In contrast, peak C at 290.2 eV, the carbonate group of 1s-π* transition,^17^ was detected from all mineral particles with more intense carbonate signal at the core region. As the mineral matured, carbonate signals became more intense and less localized, supporting the Raman analysis showing spatial distributed carbonated HAp across the interface. This variation in local chemical environment between each particle with distinct morphologies likely play key roles in mediating the assembly of CaP minerals at such an ultrathin interface region. Moreover, the polymorph features of minerals are reported to provide the most energetically favorable interaction with collagen matrix,^44^ which appear to aid the robust attachment of cartilage onto bone and toughening mechanism of the interface.

**Figure 4.**
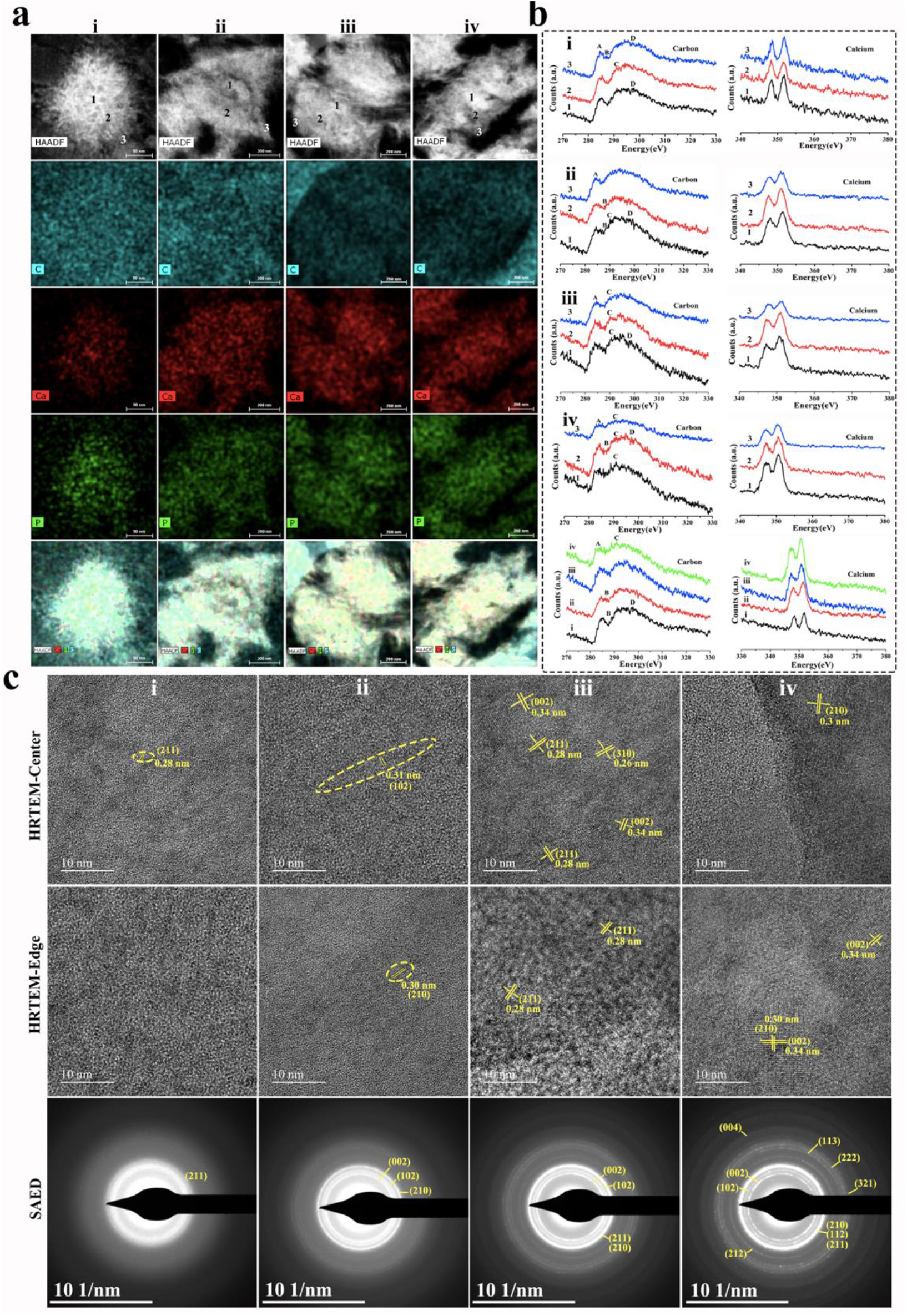
HAADF-STEM images, EDX mapping, EELS spectra, HRTEM and SAED analyses of different mineral particles in the osteochondral interface. **(a)**. HAADF-STEM images showing the characteristic mineral morphologies found in the osteochondral interface. Elemental mapping of C, Ca and P of the corresponding mineral particles. **(b)**. EELS spectra taken at C-K edge and Ca-L_2,3_ edge collected at the numbered sites (1-4) from different mineral particles (i-iv). **(c)**. HRTEM images and SAED patterns of different mineral particles at the center and edge sites, respectively. Scale bar in a for particle i = 90 nm, ii-iv = 200 nm; scale bar in HRTEM images = 10 nm; all SAED scale bars = 10 1/nm.

To further confirm how the chemical coordination environment impacted the mineral assembly and gradient transition, we performed HRTEM and SAED. HRTEM imaging and SAED demonstrated that the particles (i-iv) imbedded in interface showed increased degree of crystallinity and crystal size, which further confirmed the polycrystalline nature of the minerals (Figure 4c). Small nanocrystals, ∼2-3 nm in size, emerged in the core of spherical particles, showed the (211) plane of HAp, while its periphery was abundant in amorphous materials. For the triangular particle, a nanorod HAp with (102) plane around 3-4 nm in width and ∼20 nm in length and a nanocrystal of 3-4 nm in diameter were detected at the core and exterior of the particle, respectively, suggesting the improved crystallinity and transformed mineral shapes to elongated morphology. Particle iii-iv were highly ordered HAp, which was assembled from smaller grains with apparent boundary and embedded continuously in the amorphous region that should be the organic matrix, producing a smooth transition between mineral and organic components. The amorphous organic components guide the pathway of mineral crystal assembly in a controlled manner (including local chemical environment like carbonate substitution), to kinetically stabilize crystal lattice and consequently form a nanoscale gradient interface to lower high-energy of crystal facets that benefits mechanical properties of interface tissue.^45–47^

At the osteochondral interface, nanometer sized mineral crystals carried most of the load due to their insensitivity to the stress concentration at flaws, whereas the flaws-the soft organic components transferred load by shear stress between minerals, thus dissipate energy to ensure the crack-resist behaviors and toughing mechanism.^48, 49^ Nanoparticles at the interface region with increased aspect ratios could also provide stepwise stiffening effect.^49^ This nanoscale heterogeneity of HAp in terms of size, shape, crystallinity, and composition across the osteochondral interface not only sustain the integrity of tissue appropriately,^26^ but also drive the superior two-stages mechanical resilience *via* promoting energy dissipation and preventing the crack propagation.^25^

### Enrichment of Giant Protein Titin within the Osteochondral Interface Tissue

In addition to the transition in two-layered micro-nano structures and mineral crystals contributed to the mechanical responses at the interface, a biomolecular difference at the interface further dictates the characteristic structural and functional features from the molecular view. Confocal images revealed region-dependent distribution of collagen-I and collagen-II at the interface (Data Figure S8). Collagen-II and collagen-I were highly abundant in AC and SB region, respectively. Compared with AC and SB tissues, they were relatively sparse in transitional interface tissues. Collagen-II distributed near AC region could be responsible for the porous structure of the stage 1, while collagen-I mediated the densely packed structure of stage 2 near SB region.^1, 50^ These results also reflected the two-layered micro-nano structure variations of the interface.

To identify the distinct proteins enriched within the interface region, liquid chromatography tandem mass spectrometry (LC-MS/MS) was further performed with the sectioned tissue slices (Figure 5a, see Supplementary file 1). Compared with AC, only one protein was significantly highly expressed in interface region. However, when compared with SB tissue, 713 of proteins were differentially expressed in the interface region; of these, 195 of proteins were more abundant in interface tissue (Figure 5b, Supplementary file 2). These results indicated interface tissue showed similarity with cartilage tissues, while expressed more distinct proteins from SB tissues. The results might be limited by the incomplete extraction of interface tissue due to its ultrathin features, which possibly leads to underestimation of the protein identification.

**Figure 5.**
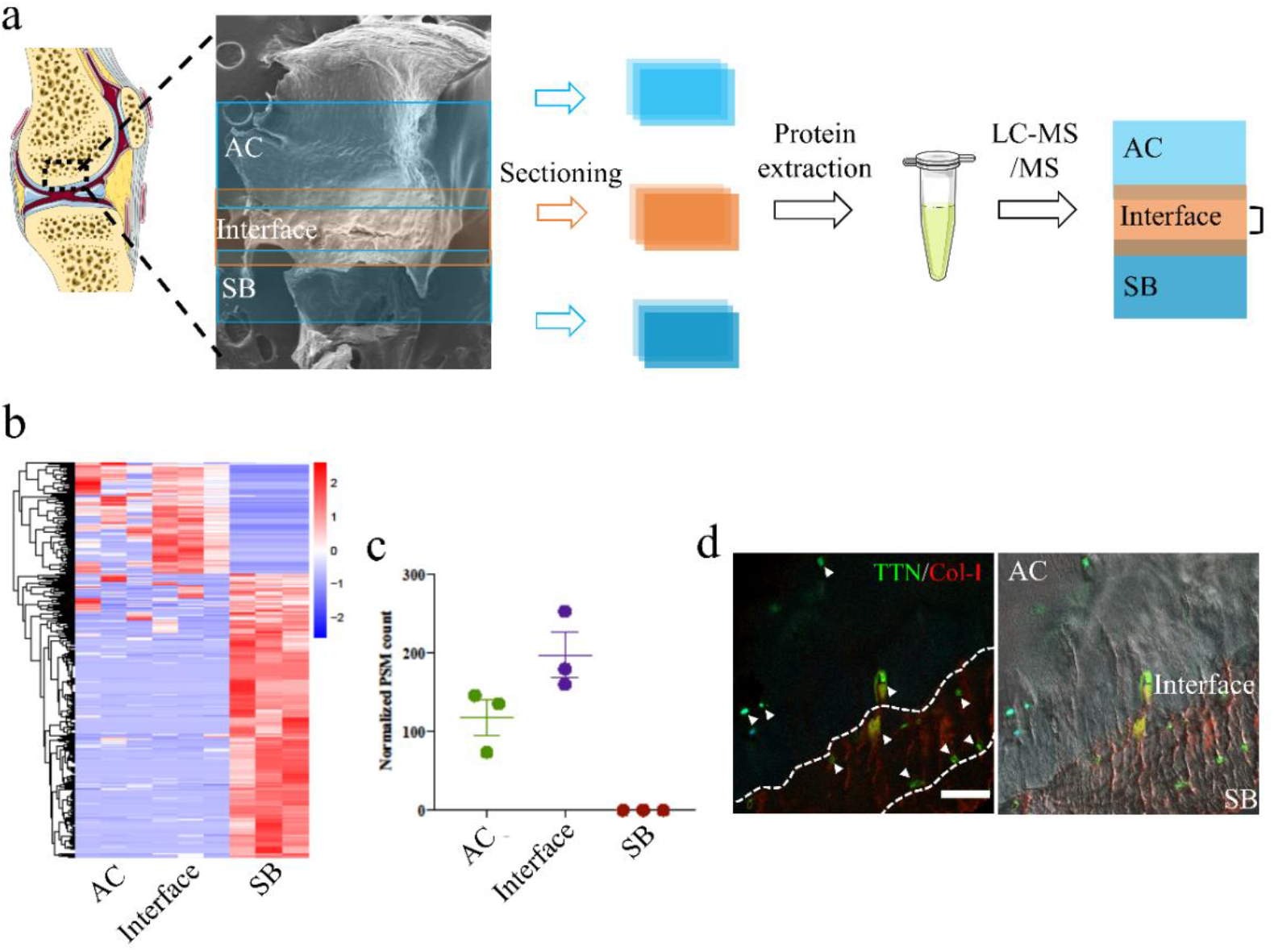
Quantitative proteome analysis of difference in expression of osteochondral interface compared to AC and SB. **(a)**. Experimental procedure for the proteomic investigation. **(b)**. Heatmap of differentially expressed proteins between osteochondral interface and SB tissue (n=3 for each tissue). **(c)**. Expression of TTN in different tissues. TTN was enriched in AC and interface region, but rarely expressed in SB tissue. **(d).** Immunofluorescence staining of TTN (green) and collagen-I (red) in osteochondral interface tissues. The osteochondral interface was clear beneath AC (collagen-I was expressed from the mineralized interface to SB region, right panel in d), and TTN was highly expressed in AC and osteochondral interface region. White arrows marked the TTN expression. White dotted lines (left panel in d) highlighted interface region. Scale bar: 20 μm.

Interestingly, compared with SB tissues, we found that giant protein titin (TTN), responsible for governing stress/strain transformation in muscle tissues,^51^ was highly expressed both in cartilage and interface tissues (Figure 5c). Markus et al. have reported TTN expression in cartilage tissues,^52^ in agreement with our results. Some other non-muscle tissues have been reported to express TTN-like proteins as stress fibres,^53–55^ while, the expression of TTN in osteochondral interface tissue has not been reported previously. We further performed immunofluorescence to confirm this using human muscle tissues as the positive control (Data Figure S9). The result showed that TTN was highly expressed in the cartilage tissue and interface region, but not in SB (Figure 5d, Data Figure S10 and S11), consisting with Figure 5c. To further examine the presence and localization of TTN, immunogold labelling was carried out. In agreement with figure 5c and 5d, we have observed positive TTN labelling in AC and osteochondral interface tissues, while failed to detect it in SB tissues (see Data Figure S10 and S11). Note that labelling was localized to chondrocytes in cartilage region; whereas, at osteochondral interface region, labelling was mainly localized around HAp minerals and sometimes adjacent to collagen fibrils. Although the banded fibril was clear, we failed to identify the collagen type, as both type I and type II collagen were present at the interface region (Data Figure S8).

As cartilage and osteochondral interface tissues shared functional similarities with muscle tissues,^52^ TTN molecules localized at cartilage and osteochondral interface region may help respond to stress/strain signaling. The elastic structure of TTN may participate in the stress transmission and endow the elasticity of osteochondral interface *via* reversible structure deformations to dissipate energy. It can be another reason responsible for the excellent mechanical response at the interface. Different from the intracellular arrangement in straited muscles, we postulated that TTN at osteochondral interface involved in both loadbearing intra- and extracellular structures.^52^ TTN molecules may further help mediate mechanical stress from the underlying SB tissue under tension and maintain cartilage homeostasis, which could be disrupted by abnormal mechanical force transferred from SB that easily results in pathological degeneration of cartilage.^56^ Moreover, the great size and high resilience of TTN may be a contributor for the integrity of osteochondral tissue.^57^ Although the presence of TTN molecules at osteochondral interface was examined, their detailed local distribution and molecular interactions with the surrounding collagen matrix and minerals were limited here. Meanwhile, deep understanding about how fundamental effects of TTN molecules on biomechanics of osteochondral interface needs scrutinizing. Future studies by employing a knockout mice model will provide insights into structures and functions of TTN in this soft-to-hard interface.

Gene ontology analysis on the proteins highly expressed in osteochondral interface was performed when compared with SB tissues. The results revealed that muscle tissue morphogenesis, blood vessel endothelial cell differentiation as well as the positive regulation of BMP signaling pathway were over-represented (Data Figure S12). Considering mineralization related proteins are also enriched in SB tissues, it is expected that there is no significant differential expression of these proteins between interface and SB tissues. These biological processes might be associated with the observed compositional changes and functional features.

Collectively, these observations suggested that osteochondral interface tissues showed superior mechanical response driven by characteristic biomolecular compositions and well-organized nanoparticle assemblies. Relying on brittle minerals and weak organic matrix as building motifs, the load-bearing transition interface plays with sophisticated design strategies at micro and nano length scale to render an excellent interface property.^58, 59^ Mineral gradients, in terms of density, morphology and crystallinity across the transverse direction of osteochondral interface, are important material motifs here used to build smooth transitional bridges between AC and SB tissues, and play curial role in regulating mechanical response.^32^ The exponential profile of interface tissues, together with porosity and mineral crystallinity differences, are considered to be the key factors in arresting cracks between the mechanically dissimilar materials.^31, 60^ FEA results further validate the superiority of exponential profile of tissue modulus in facilitating force transfer. The nanoscale heterogeneity of minerals also contributes to the toughing mechanism of interface tissues.^25, 49^ Changes of local chemical environment, especially for decrease of carbonate concentrations along longitudinal direction, largely affect morphologies and crystal crystallinity of HAp, resulting in mediated tissue modulus for interface tissue.^42^ Interestingly, elastic protein-TTN, mainly expressed in straited muscles, is detected to be enriched in osteochondral interface tissues, suggesting to elevate tissue elasticity and may help dissipate energy via reversible deformation and damp forces when joint move.^61^

Taken together, the two-layered micro-nano structures, sophisticated HAp assemblies, molecular composition and exponential increment of tissue modulus enable the interface tissue to withstand compress stresses and ensure integrity of interface tissues. The graded toughing composite materials enable effective force transfer and avoid stress concentrations localized in interfaces between mechanically dissimilar materials.

## CONCLUSIONS

The osteochondral interface tissue was comprehensively studied and exhibited a two-layered ultrathin calcified region of 20-30 μm where porous structure of collage II was gradually changed to be dense structure with densely mineralized nanocomposites within collagen I (Figure 6). This gradient localized carbonated HAp nanoparticles assembled within organic matrix along longitudinal direction allowed the resilient attachment of cartilage onto bone and built a natural barrier between them to maintain their respective functions. The mechanical properties of the interface tissue showed an exponential increase (∼3 orders of magnitude) of elastic modulus, which was proved by FEA simulation to be rational to facilitate force transfer and reduction of stress concentration, thus avoiding tissue failure. The nanoscale heterogeneity of carbonated HAp in terms of shape, crystallinity as well as compositions in the transverse direction of interface played a key role in dissipating energy and increasing toughness during joint movement. Strikingly, the giant protein titin enriched in the interface seemed to endow the tissue with elasticity and help the tissue counteract compressive stress under tension condition. By combining characterization of microstructural, micromechanical, nano-compositional and biomolecular features of the interface, we clearly elucidated the mechanism of the unusual fatigue-resistant adhesion and toughening properties of cartilage-to-bone interface tissue. The identified mechanism of soft-to-hard interface that allows effective force transfer at certain direction may provide potential guideline for future generation of biocomposite interface materials design.

**Figure 6.**
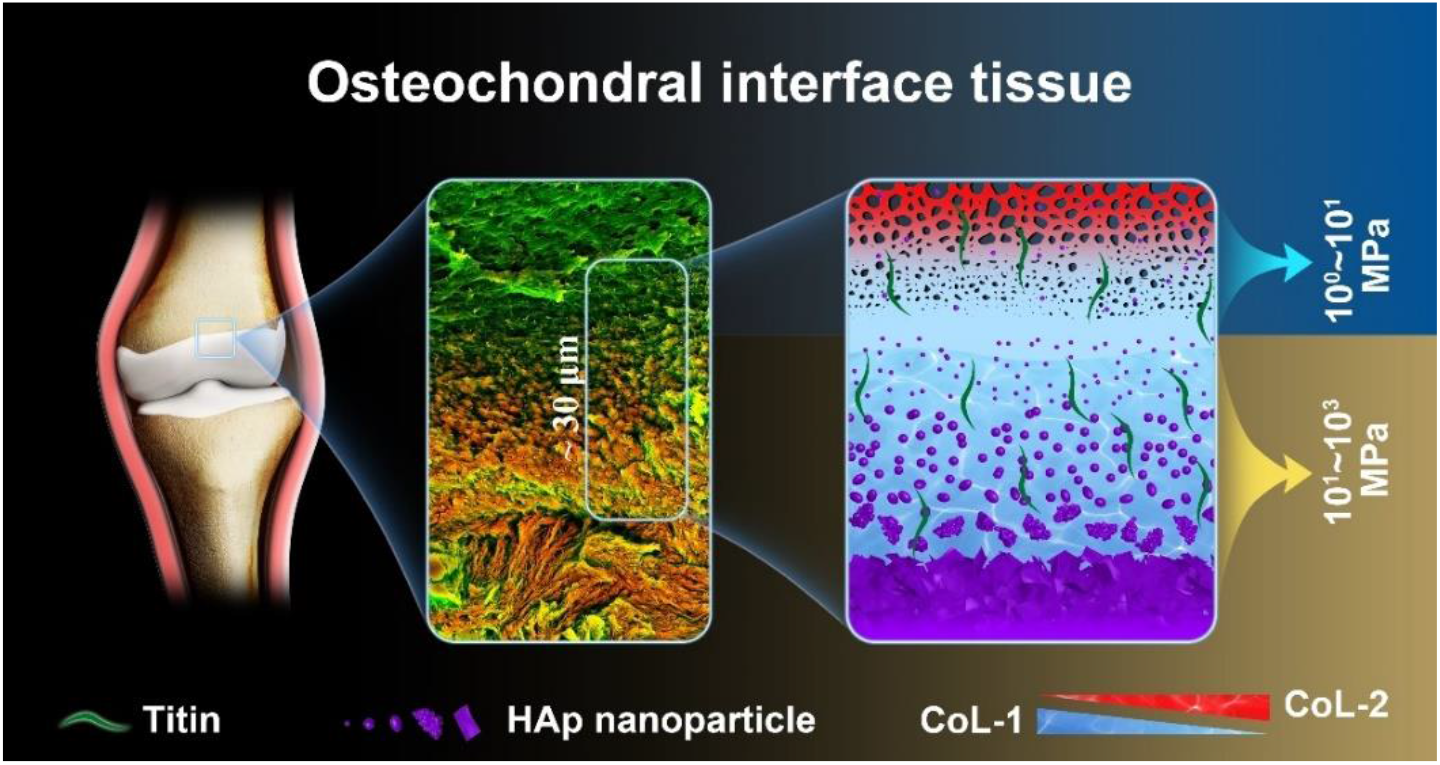
Schematic showing the specific two-layered micro-nano structure variation, nanocomposite assembly and biomolecular components of the ultrathin osteochondral interface. Within 20∼30 μm, the interface tissue underwent a characteristic two-layered transition in microstructure, nano-HAp assembly and compositions. Before attaching to the SB tissue, porous structure gradually changed to be densely packed materials, nano-HAp varied in terms of morphology, crystallinity and chemical compositions, and elastic-responsive protein-TTN was enriched in the interface tissue. These variations together contributed to the excellent mechanical properties (three orders increment of magnitude for tissue modulus) of the ultrathin osteochondral interface.

## EXPERIMENTAL SECTION/METHODS

### Sample Preparation

All normal human osteochondral interface samples were collected as surgical waste under ethics approval from Second Hospital of Shanxi Medical University Ethics Committee of four male individuals (26, 28, 43 and 48 years old) with informed consent from patients. Healthy human knee joints were obtained at the time of surgical amputation. Tissues were dissected into small pieces and washed with sterile phosphate buffered saline (PBS) before storage at −80 °C when not undergoing treatment and analysis.

### Histology

Normal osteochondral sample slices were fixed in 4% paraformaldehyde for 48 h followed by cryo-sectioning into 15-μm sections for Safranin-O (SO) staining and cryo-sectioning into 200-μm sections for electron microscopy, respectively. Samples of 15-um thick were stained with SO according to standard protocol to detect cartilage components and the osteochondral interface.

To detect the organic matrix interaction of osteochondral interface, samples were fixed in 4% paraformaldehyde, decalcified in 10% EDTA-2Na (w/v) solution and cryo-sectioned into 200-μm thick, followed by stored at −80 °C for further analysis.

### Immunofluorescent Staining

The frozen cryocut sections were first thawed at room temperature and washed with PBS solution (pH 7.2) to remove the O.C.T compound. After the heat-induced antigen retrieval process using sodium citrate (10 mM) at 65 °C overnight, samples were blocked with 5% (wt/v) Bovine Serum Albumin (BSA, Sangon Biotech, China) for 60 min at room temperature. Then, samples were incubated with different primary antibodies for Titin antibody (1:100, bs-9861R-AF350, Bioss), Anti-Collagen-I antibody (1:200, ab6308, Abcam; and 1:200, Catalog No.AF7001, Affinity) and Anti-Collagen-II antibody (1:200, ab185430, Abcam) at 4 °C overnight. Then, samples were washed with PBS-Tween (0.01% Tween) and incubated with corresponding secondary antibodies (1:200 diluted in PBS) at room temperature for 90 min: Alexa Fluor® 488 goat anti-rabbit IgG antibody (Product # A-11008, Invitrogen); Donkey anti-Mouse IgG Secondary Antibody, Alexa Fluor 546 conjugate (Product # A10036, Invitrogen). After rinsing with PBS solution, samples were mounted with Fluorescence Mounting Medium for further imaging by using a laser confocal microscope (Zeiss 880). The acquired images were analyzed with Image J.

### Immunogold labelling

The fresh cartilage tissues (∼1×3 mm) were first fixed in a 3% (w/v) formaldehyde + 0.1% glutaraldehyde (w/v) in PBS solution at 4 °C for 3∼4 h, then dehydrated through a graded ethanol treatment (30 and 50 % (v/v) for 30 min at 4 °C, 50, 70, 90, 100, 100, 100 % (v/v) for 1 h at −20 °C in each solution). Then, samples were infiltrated with Lowicryl resin (Polysciences, Inc., Warrington, PA) series diluted with ethanol at 30, 70 and 100 % for 2 h each at −20 °C, and then soaked with pure resin overnight at −20 °C. Finally, the resin solution was changed and the samples were curetted by ultraviolet light at −20 °C for 3 days and at room temperature for 2 days. The curetted resin blocks were then sectioned at 100 ∼ 150 nm and adsorbed to carbon-coated nickel grids. The grids were floated with 50 μL PBS and deionized water for 5 min, and then blocked in 1 % BSA (w/v) + 0.05 % Triton X-100 (v/v) +0.05 % Tween 20 (v/v) in PBS solution for 30 min at room temperature. After that, the grids were floated in a 30 μL drop of primary antibody diluted with the blocking agent (anti-titin antibody, 1:50, bs-9861R-AF350, Bioss) overnight. The grids were washed with PBS and deionized water 5 times for 2 min each, and then floated in 30 μL drops of immunogold solution (10 nm gold-anti-rabbit, Jackson ImmunoResearch Laboratories, West Grove, PA, concentration for 1:30) for 1 h. Grids were washed and air dried, and stained with 2% uranyl acetate and finally imaged with TEM at 80 kV (Hitachi 7650).

### Synchrotron Radiation-Microcomputed Tomography

The microstructure of osteochondral interface was evaluated by synchrotron radiation-microcomputed tomography (SR-μCT) at the BL13W1 of Shanghai Synchrotron Radiation Facility in China. The samples were scanned at a beam energy of 18 keV with exposure time of 0.5 s. Sample-to-detector distance was 5.0 cm. For each acquisition, an angular step of 0.15° over an angular range of 180° was used to capture 1200 projection images by the CCD detector with a pixel size of 3.25 μm. Dark-field and flat-field images were also collected to correct electronic noise and reduce the ring artifact during reconstruction. These projected images were sequentially phase-retrieved, transformed into 8-bit slices by the software of PITRE written by BL13W1, and reconstructed to acquire 3D images using Image Pro Analyzer 3D software.

### SEM and EDX Analyses

200-um-thick cryo-cut samples were thawed at room temperature and washed with DI water for several times, followed by dehydrated through a graded ethanol treatments (20, 30, 40, 50, 70, 80, 90, 95, 100 and 100 % (v/v)) for 30 min in each solution and air dried. Then, the samples were gold sputtered and imaging was performed (Hitachi SU-70) at 3 kV accelerating voltage to observe the morphology of osteochondral interface. Energy dispersive X-ray spectroscopy (EDX) line scan were performed to identify the elements distribution across the interface. As for the decalcified samples, prior to dehydrated treatments and SEM, the cryo-sectioned slices were firstly immersed in 10% (w/v) EDTA-2Na solution for at least 7 days to remove the minerals.

### AFM

Surface topography features of cartilage-to-bone interface were detected by atomic force microscopy (AFM, NTEGRA Spectra, NTMDT) with a tapping mode.

### Nano-FTIR

The chemical composition of osteochondral interface was detected using a Nano-IR spectrometer equipped with a broadband DFG laser for spectroscopy and a tunable mid-IR laser (QCL) for IR imaging (Anasys Instruments, nanoIR2-fs). Each spectrum was the result of signal-averaging of 64 scans with the wavenumber ranging from 400 to 4000 cm^−1^.

### XRD

The crystalline structure of the mineralized interface between cartilage and bone was measured using X-ray powder diffraction (X’Pert^3^ Powder, Malvern Panalytical, Netherlands) with Cu Kα radiation (λ= 0.154 nm) operating at 40 kV. The data were scanned with a sampling step of 0.2° in the 2θ range from 5° to 70°.

### Micro-Raman Spectroscopy

The Micro-Raman spectroscopy was performed to measure the mineralized components and their distributions of osteochondral interface using a confocal Raman microscope (LabRAM HR Evolution, Horiba Co., Ltd.) equipped with a 532 nm laser. Spectra were collected in the 300∼2000 cm^−1^ using an electron multiplying charge coupled device (EMCCD) detector at a spectral resolution of ∼2 cm^−1^. Raman images (50 × 50 um, spatial resolution ∼2 μm) of osteochondral interface were obtained by continuous scanning. Each spectrum was collected with an acquisition of 1 s and 1 accumulation. No sample degradation was noticed using this parameter.

### Nanoindentation

The mechanical properties of osteochondral interface region were tested using a nano-indenter to measure the surface stiffness of tissue (PIUMA, Optics 11, Amsterdam, The Netherlands). A soft cantilever with a radius of 50 μm and a stiffness of 267 N/m were used for the measurement at an indentation depth of 3000 nm. A nanoindentation test matrix comprising 20×20 indents were performed on osteochondral tissue region from articular cartilage (AC) to subchondral bone.^62^ Distance between two adjacent locations were 10 μm. The efficient Young’s modulus was calculated from the slope of the unloading curve by using a power law fit to 20∼90% of the unloading curve with Hertz model. Three different specimens were tested.

### Cryo-SEM

In order to observe the microstructure of osteochondral interface *in situ*, Cryo-SEM imaging was performed on a FEI Helios NanoLab 600i Analytical Field Emission Scanning Electron Microscope fitted with low temperature sample carrier Quaroum PP3000T. The sample was loaded on the cryo-specimen holder and cryo-fixed in slush nitrogen (−210 °C), followed by being transferred into the cryo-stage. Then, the sample was fractured to get a fresh interface. The temperature of the sample was raised by heating the holder to −90 °C for 60 min, in order to increase the contrast and sublimate free water in the solid-state lakes, followed by a temperature decrease to −180 °C, to stabilize the sample. The surface of the frozen preparation was then coated with platinum (10 mA, 30 s) to prevent charging of the sample and to obtain a good relation between signal and noise. The coated sample was thereafter transferred into the microscope chamber where it was analyzed at a temperature of −180 °C.

### TEM, EDX, EELS and SAED analyses

The specimens were fixed in 2.5% (v/v) glutaraldehyde in phosphate buffer (0.1M, pH: 7.2) for 24 h, washed several times with PBS solution, bulk stained with 1% (v/v) OsO4 in 0.1 M phosphate buffer, dehydrated, and embedded in a spurr-based resin. Then, the specimens were cross sectioned in Lerca EM UC7 ultratome at 100 nm onto bare 300-mesh nickel grids. Preliminary bright-field TEM imaging and EDX of the cross-sectioned osteochondral interface was conducted on a field-emission FEI Tecnai G2 F20 microscope (FEI) operated at 200 kV. HAADF-STEM, STEM-EDX mapping, SAED and STEM-EELS were performed with an aberration-corrected scanning transmission electron microscope (FEI Titan G2 80-200 microscope equipped with a Super-X EDX detector) operated at 200 kV and 300 kV, respectively. The microscope was equipped with a DCOR plus spherical aberration corrector for the electron probe which was aligned before every experiment by using a gold standard sample. Calcium-to-phosphorus (Ca/P) ratios of different zones in the interface were calculated. EELS spectrum images were collected from different region of interest in the range of 250∼400 eV to investigate the fine structure of C (K-edge) and Ca (L_2,3_-edge). Spectra were calibrated, normalized, and background subtracted and processed by principal component analysis (PCA). No obvious damage was observed during the experiments. Gatan DigitalMicrograph software was used for STEM image, SAED and EELS analysis.

### Proteomics

Sample preparation: AC, osteochondral interface and SB tissue (n=3) were prepared by cryo-sectioning. it’s noted that the difference between interface, AC and SB were mostly larger than our results here since the interface samples were not fully distinguished from AC and SB without labelling. AC and SB tissues were prone to remained in the interface region. Three kinds of tissue were collected in the tube and cut into small pieces. After being washed with PBS, the tissues were grinding through pestles under liquid nitrogen condition until it became power, cells were lysed in cold modified RIPA buffer (50 mm Tris-HCl, pH 7.5, 1 mm EDTA, 150 mm NaCl, 1% N-octylglycoside, 0.1% sodium deoxycholate, complete protease inhibitor mixture) and incubated for 30 min on ice. After that, lysates were cleared by centrifugation. 200 μg protein samples from the tissues were counted by BCA assay. Then, the protein samples were reduced by DTT and alkylated by iodoacetamide. All crude protein extracts were precipitated by acetone precipitation at −20 °C for 2 h, then the precipitated protein samples were re-dissolved in 200 uL TEAB buffer. Finally, trypsin was added (trypsin/protein, 1:50), and the solution incubated at 37 °C for 12-16 h. Digestion was stopped by the addition of 2% trifluoracetic acid. The tryptic digests were desalted with C18 solid-phase cartridges. Peptides were separated by HPLC focusing into 9 fractions.

Reverse-phase high-performance liquid chromatography and mass spectrometry (RP-HPLC): RP-HPLC was performed. Briefly, the first dimension RP separation by micro-LC was performed on a U3000 HPLC System (Thermo Fisher Scientific, USA) by using a BEH RP C18 column (5 μm, 300 Å, 250 mm × 4.6 mm i.d., Waters Corporation, USA). Mobile phases A (2% acetonitrile, adjusted pH to 10.0 using NH_3_·H_2_O) and B (98% acetonitrile, adjusted pH to 10.0 using NH_3_·H_2_O) were used to develop a gradient. The solvent gradient was set as follows: 5-8% B, 1 min; 8-50% B, 6 min; 50-80% B, 14 min; 80-95% B, 1.5 min; 95% B, 7 min; 95-5% B, 2 min; 5% B 5 min. The tryptic peptides were separated at an eluent flow rate of 1.0 mL/min and monitored at 214 nm. The column oven was set as 45 °C. Eluent was collected every 90 s. Forty fractions were collected for each sample. The samples were dried under vacuum and reconstituted in 15 μL of 0.1% (v/v) FAin water for subsequent analyses. Fractions from the first dimension RPLC were dissolved with loading buffer and then separated by a C18 column (75 μm inner-diameter, 360 μm outer-diameter × 15 cm, 2 μm C18). Mobile phase A consisted of 0.1% formic acid in water solution, and mobile phase B consisted of 0.1% formic acid in 80% acetonitrile solution; a series of adjusted linear gradients according to the hydrophobicity of fractions eluted in 1D LC with a flow rate of 300 nL/min was applied. The MS conditions were as the followings: For Orbitrap Fusion Lumos, the source was operated at 1.9 kV, with no sheath gas flow and with the ion transfer tube at 350 °C. The mass spectrometer was programmed to acquire in a data dependent mode. The survey scan was from m/z 350 to 1500 with resolution 60,000 at m/z 200. The 20 most intense peaks with charge state 2 and above were acquired with collision induced dissociation with normalized collision energy of 30%, one microscan and the intensity threshold was set at 1000. The MS2 spectra were acquired with 15, 000 resolution. The peptides were detected, isolated, and fragmented to produce a tandem mass spectrum of specific fragment ions for each peptide.

Mass spectrometry data analysis: MS peptide sequences and protein identity were determined by matching fragmentation patterns in protein databases using the Mascot software program (Matrix Science, Boston, MA). Enzyme specificity was set to partially tryptic with two missed cleavages. Modifications of the peptides included carboxyamidomethyl (cysteines, variable), oxidation (methionine, variable), phosphorylation (S, T, Y, H, variable) and acetylation (N-term, K, variable). Mass tolerance was set to 20 ppm for precursor ions and fragment ions. The database searched was Uniprot (Homo sapiens).

Protemoics data analysis: The differentially expressed proteins were identified by limma R package with peptide-spectrum matches (PSMs) information as input.^63, 64^ The data was log2-transformed and normalized *via* the normalizeBetweenArrays function. Then, the lmFit function was used to fit a linear model for each protein. The eBayes moderated t-statistic was calculated for each protein and p-values were corrected for multiple testing by the positive false discovery rate (FDR) *via* the contrasts.fit and eBayes functions. Differentially expressed proteins were defined as differential expression of >2 fold and adjusted p value < 0.05.

### Theoretical Model

In order to explore the function of the spatial distribution of the modulus at the osteochondral interface, we proposed a simple theoretical model to demonstrate its superiority. Based on the consideration that the osteochondral interfaces were currently placed under a quasi-static compression loading,^65^ a two-dimensional shell suffering uniaxial compression was employed to evaluate the function of gradient modulus at the interface. The length and height of the shell were 10 μm, respectively. Three gradient forms of the elastic moduli along the transverse direction were presented in our study, which were linear (*E*(*y*)=2.21×10^7^*y*-1.81×10^7^), power (*E*(*y*)=4×10^6^*y*^1.367^), and exponential (*E*(*y*)=2×10^6^*e*^0.069y^), respectively.^35^ Since the exponential distribution was the closest form to the experiment data, the comparison of the mechanical behavior among different distribution forms could reveal the excellence of such a particular modulus distribution sufficiently.

### Finite Element Analysis

Finite element analysis (FEA, Abaqus/CAE 6.13) was used to simulate the model. The model was divided into 100 layers for discretization with assuming the modulus in each layer was identical. The interface model was meshed using Plane 8-node biquadratic shell elements and assigned linear elastic material properties. The number of elements and nodes were 10000 and 30401, respectively. The elastic modulus of each layer in the simulation was taken from the model function we set and the Poisson’s ratio was taken 0.3 for all layers^28^. Meanwhile, the load for compression displacement loading is set as 0.1μm. The von Mises stress was selected to measure the mechanical behavior of the interface region which was defined as:

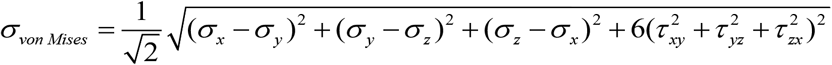

### Statistical Analysis

All data are presented as the mean ± SD. Differences between the values were evaluated using one-way ANOVA analysis (Tukey’s post-hoc test). p < 0.05 was considered statistically significant.

## ASSOCIATED CONTENT

### Supporting Information

The supporting information available free of charge at https://

The gross appearance of human knee joint samples, SO staining images, AFM images, FTIR spectra, SEM and EDX spectra, XRD diffraction patterns, FEA displacement loading results, tables with approximate transitions, confocal images, gene ontology enrichment related to biological process on the upregulated proteins of osteochondral interface tissues

## Author Contributions

X.W. and J.L. contributed equally. X.W., J.L. and H.O. conceived the project; X.W., J.L., Y.M., X.Z., Q.W. and H.Q. carried out the experiments; Z.L. performed the FEA simulation; J.L. and S.X. performed the proteomics work; X.W., Z.L. and H.O. wrote the manuscript; all authors commented on the manuscript.

## Notes

The authors declare no competing financial interest.

## ACKNOWLEDGMENT

We thank Mr Nianhang Rong for the SEM imaging and EDX analysis; we thank Miss Guoqing Zhu for the HRTEM and STEM imaging and EDX analysis.

