## Supplementary material for "High Resolution Nanostructure with Two-stages of Exponential Energy Dissipation at the Ultrathin Osteochondral Interface Tissue of Human Knee Joint": Supporting Information.docx

^⊥^China Orthopedic Regenerative Medicine Group, Hangzhou (CorMed), Hangzhou, 310058, China.

^#^Department of Engineering Mechanics, Zhejiang University, Hangzhou, 310027, China.

^∇^Key Laboratory of Structural Biology of Zhejiang Province, School of Life Sciences, Westlake University, Hangzhou, 310024, China.

^▲^Department of Orthopedics, Second Hospital of Shanxi Medical University, Shanxi Key Laboratory of Bone and Soft Tissue Injury Repair, Taiyuan, 030001, China.

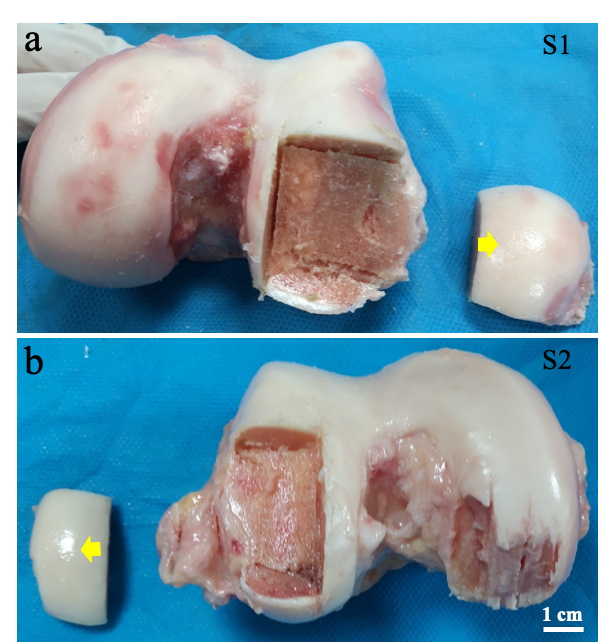

**Figure S1. Gross appearance of human knee joint samples.** S1 sample was obtained from the knee of a 43 years old male (a), and S2 sample was from a 26 years old male at the time of surgery (b). The arrowheads highlighted the cartilage tissue we used. Scale bar: 1 cm. (Photo credit: Xiaozhao Wang, Zhejiang University)

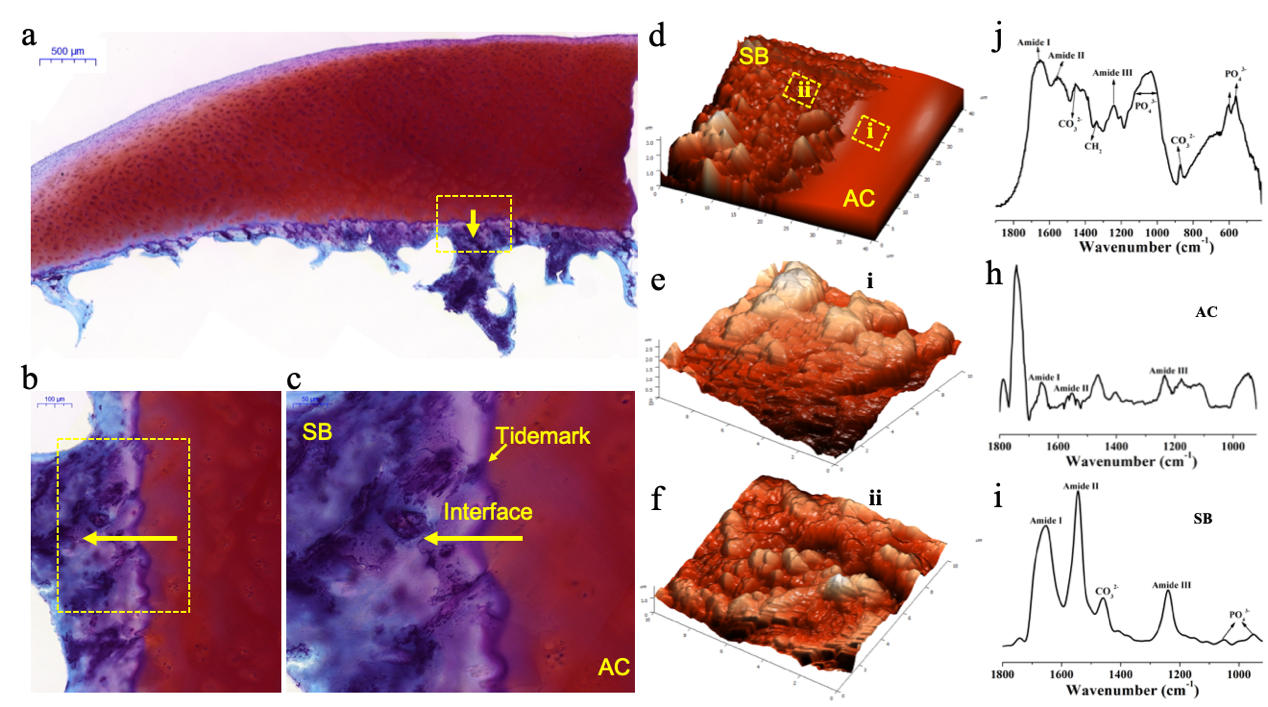

**Figure S2.** **(a-c)**. Safranin O/Fast Green (SO) staining of the articular cartilage tissue for S2 sample. The arrow indicated the tidemark of the tissue (c). The boxed area showed the transitional area between the articular cartilage (AC) and subchondral bone (b). **(d)**. AFM topography image of the osteochondral interface. **(e)** and **(f)** were the enlarged images of AC area and SB area. **(j-i)**. FTIR results of osteochondral interfaces, AC and SB region, respectively, corresponding to the area of e-f. The transition from AC to SB exhibited obviously heterogeneous topography. AC tissues (average roughness: 72 nm) were much smoother than SB sites (average roughness: 210 nm). The FTIR spectra showed the existence of organic components, such as collagen and GAGs, in AC area, while the emergence of PO_4_^3-^ and CO_3_^2-^ revealed the mineral components in SB region, indicating the compositional transition of the osteochondral interface.

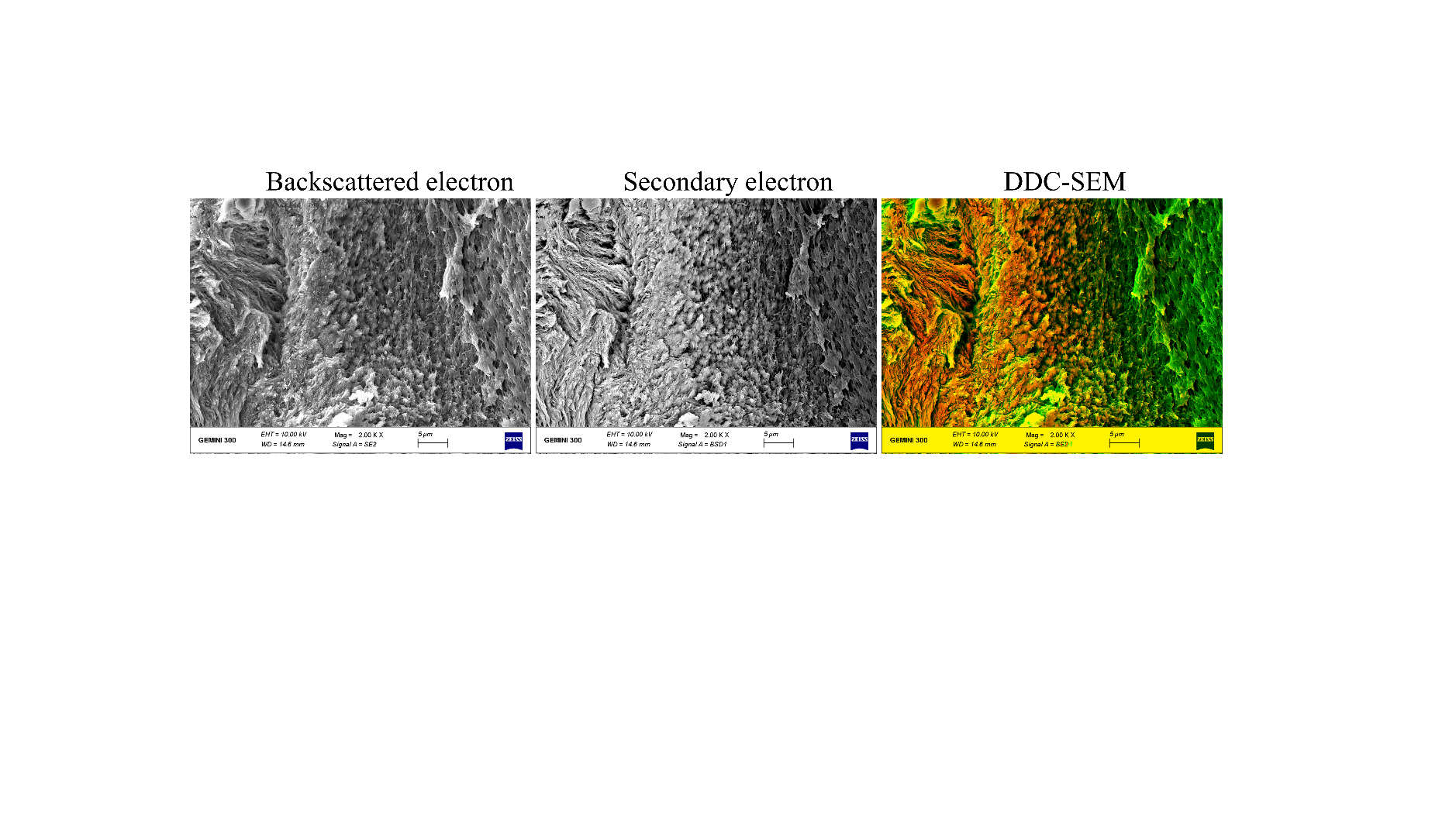

**Figure S3.** Image processing steps used to creat a density dependent colour scanning electron micrographs (DDC-SEM) showed in Figure 1d. The backscattered electron images and secondary electron images were assigned to be red and green channels, respectiely, follwed by stack command to acquire a combined images. After that, the combined images were converted to single RGD images. The final output images clearly presented mineral morphologies and distributions across the interface region.

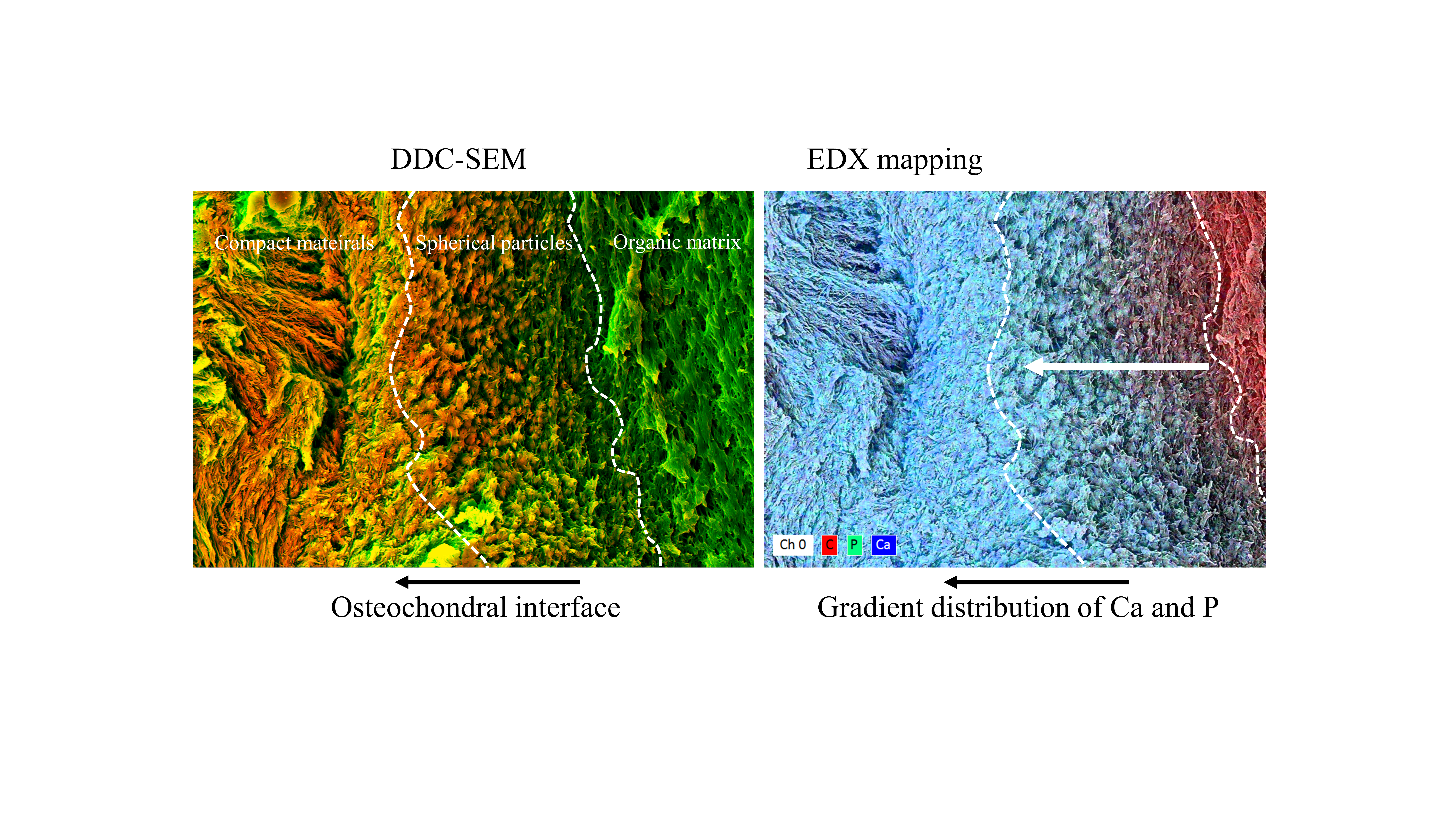

**Figure S4.** The corresponding EDX elemental maps of the sectioned osteochondral interface (right). Ca (blue), P (green), and carbon (magenta) were distributed hierarchically throughout the interface. The Ca/P-rich region was complementary to the C-rich region.

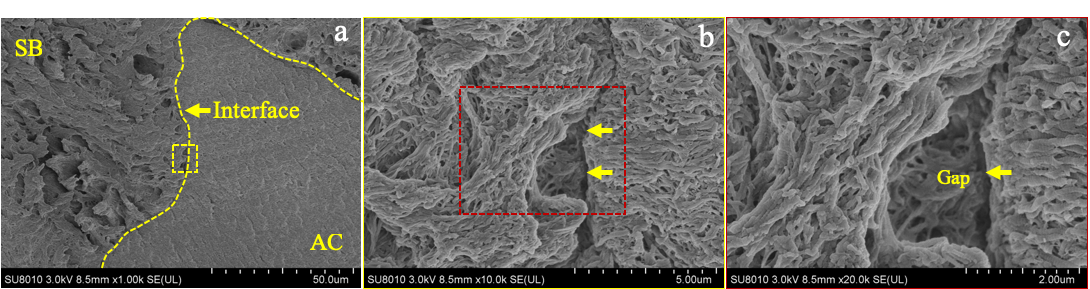

**Figure S5. (a-c)**. SEM images of the decalcified osteochondral tissue of S2 samples. After decalcification, it’s notably that collagen fibrils organization were apparently varied between AC and SB area. Also, mineral particles in the interface region were diminished and obvious gap was emerged, revealing that mineral particles played important roles in regulating the adhesion of cartilage to the bone. Additionally, the collagen arrangement and organization in the interface were significantly different from that in AC area.

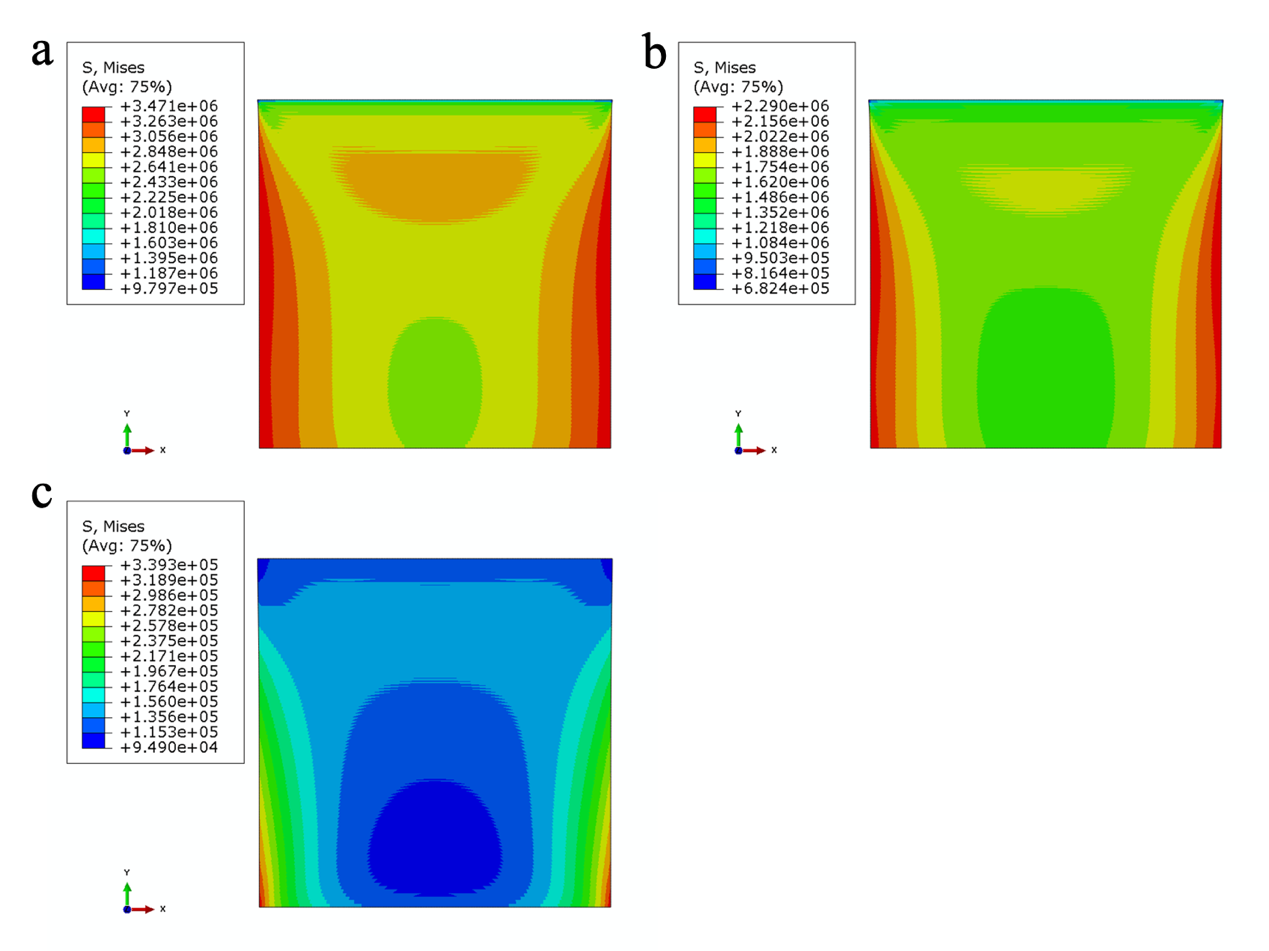

**Figure S6.** **FEA results (Displacement loading)**. Von Mises stress (Pa) distribution with the different function forms of elastic modulus. **(a)**. Linear; **(b)**. Power; **(c)**. Exponential.

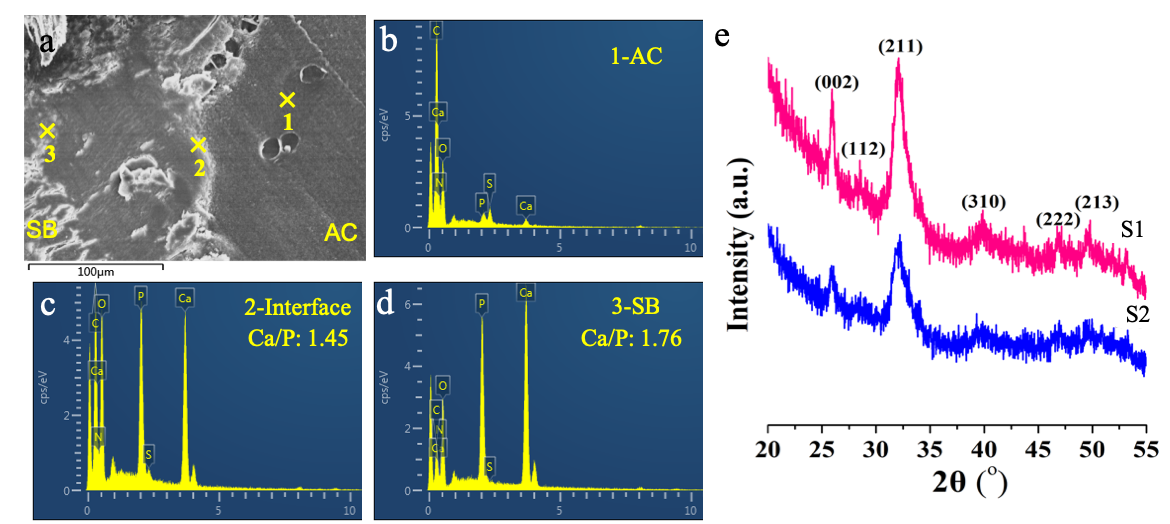

**Figure S7.** **Elemental analyses of osteochondral interface of S2 sample. (a)**. Microstructure of osteochondral interface of S2 sample, 1, 2 and 3 represented AC site, interface and SB region, respectively. **(b-d)**. corresponding EDX spectra collected at the numbered sites indicated on the micrograph a. Scale bar, 100um. The calcium to phosphorus (Ca/P) ratio of the interface was about 1.46, while that of the SB region was 1.76 (natural bone: hydroxyapatite (HAp)), indicating different crystal phase and composition of CaP in the osteochondral interface. **(e)**. XRD diffraction patterns of S1 and S2 can be indexed to HAp (JCPDS no. 09-0432) crystals, characterizing the presence of HAp in the osteochondral interface.

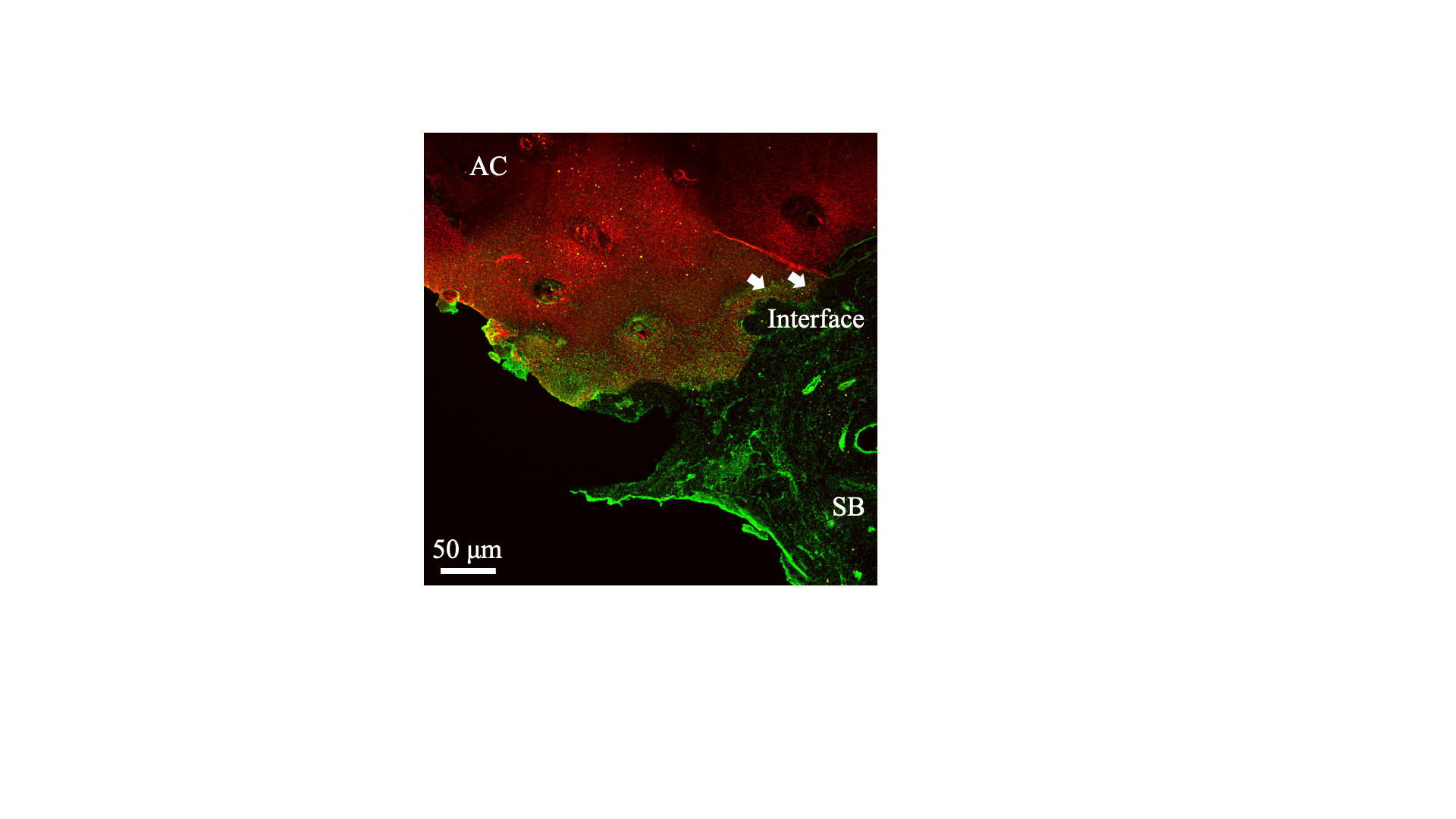

**Figure S8.** Confocal images of osteochondral interface tissue immune-stained with collagen-I (green) and collagen-II (red).

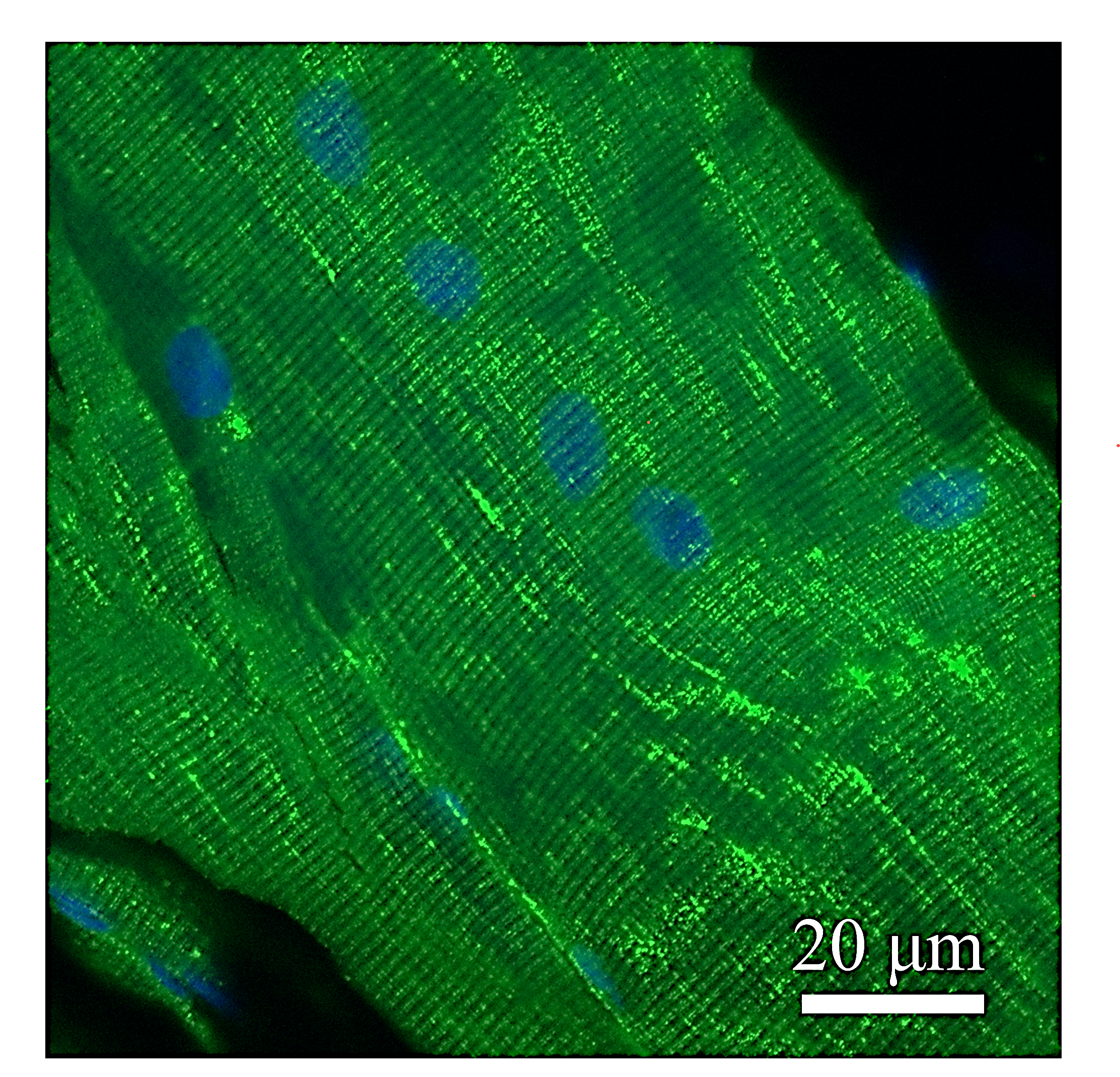

**Figure S9. Immunoflurescence staining of titin (TTN) in huaman muscle tissues.** Scale bar = 20 μm

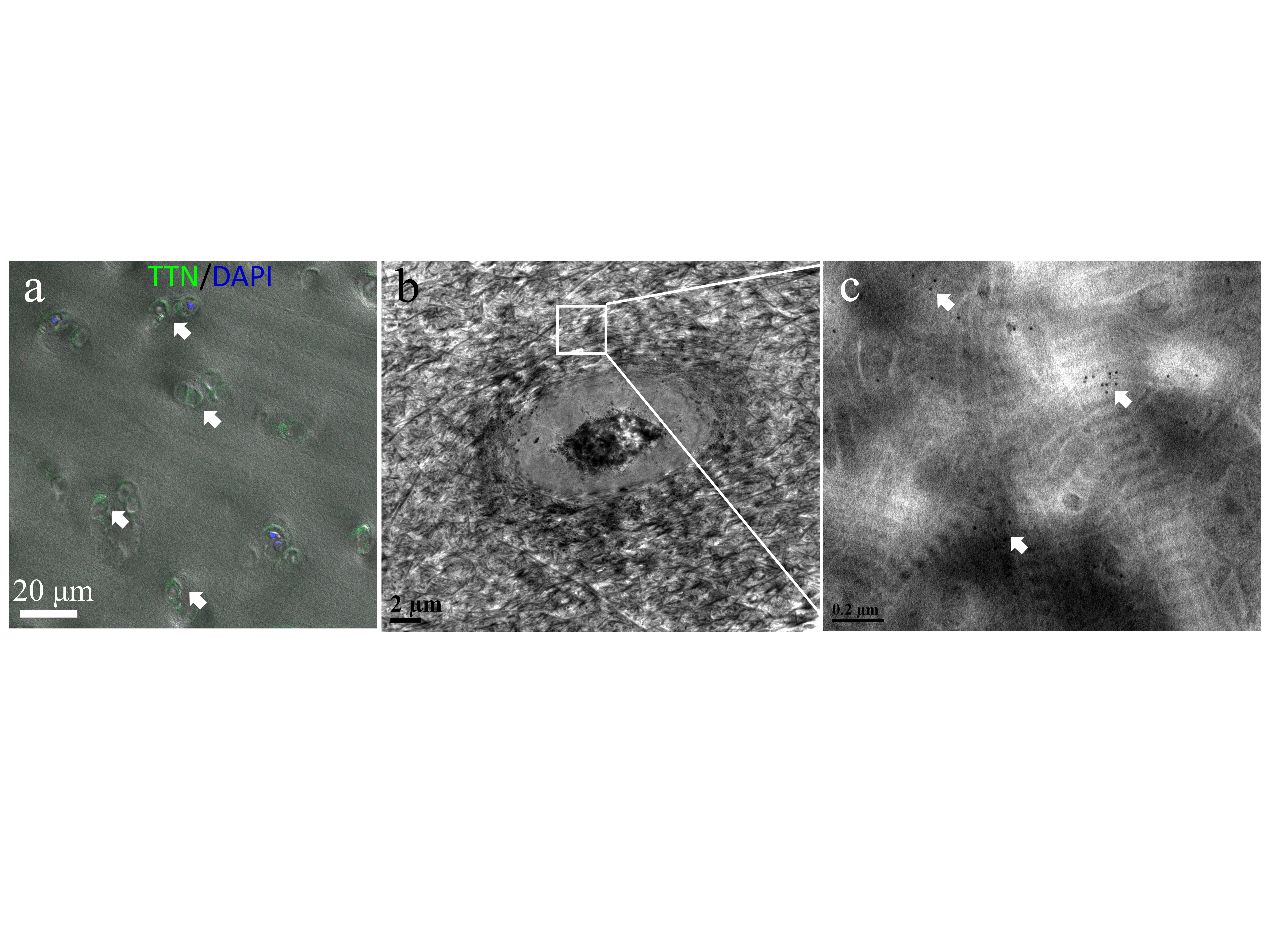

**Figure S10. TTN localization around chondrocytes at cartilage revealed by immunofluorescence staining and immunogold labelling**. **(a)**. Immunofluorescence staining of TTN (green) expressed at cartilage tissues. **(b-c)**. TEM images of cartilage tissues presenting positive immunogold labelled for TTN (white arrows: 10 nm nanoparticles). Labelling was localized to chondrocytes, which is consistent with immunofluorescence staining results. Scale bars, in a = 20 μm, b = 2 μm, c = 200 nm

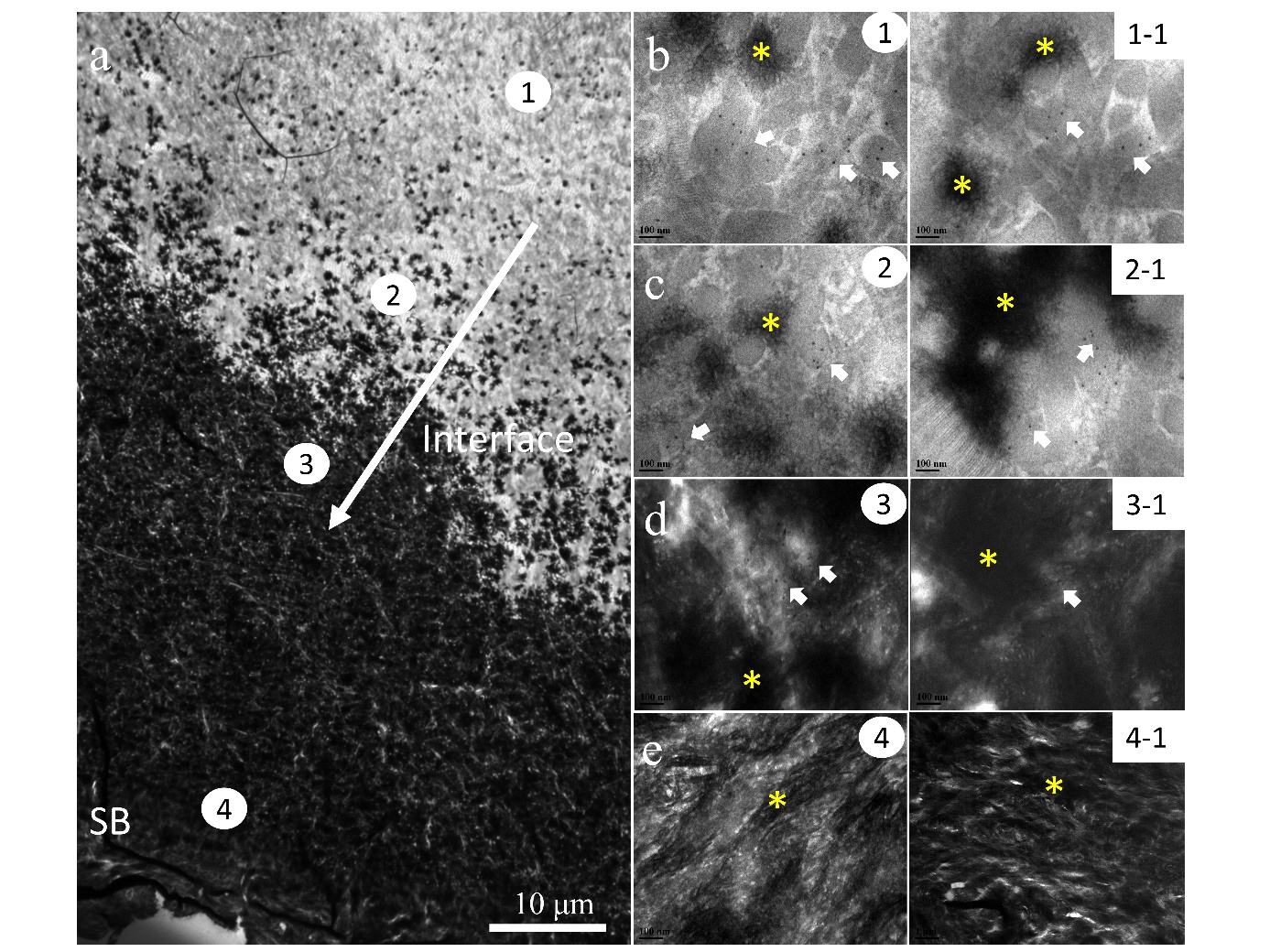

**Figure S11.** **Immunogold labelling of TTN molecules revealing their ultrastructural localization in the ECM of osteochondral interface tissues.** **(a)**. Transmission electron microscopy (TEM) images presenting gradient HAp distributions of osteochondral interface tissues. **(b-e)**. TEM images of numbered sites at osteochondral interface tissue (b-d) presenting positive immunogold labelling for TTN, but it was difficult to identify labelled gold at SB tissues (e). Gold nanoparticles were evident and highlighted with white arrows. HAp particles were marked with yellow asterisk. Note that labelling was localized around HAp particles and were sometimes adjacent to collagen fibrils. Scale bars in a = 10 μm, b-e = 100 nm.

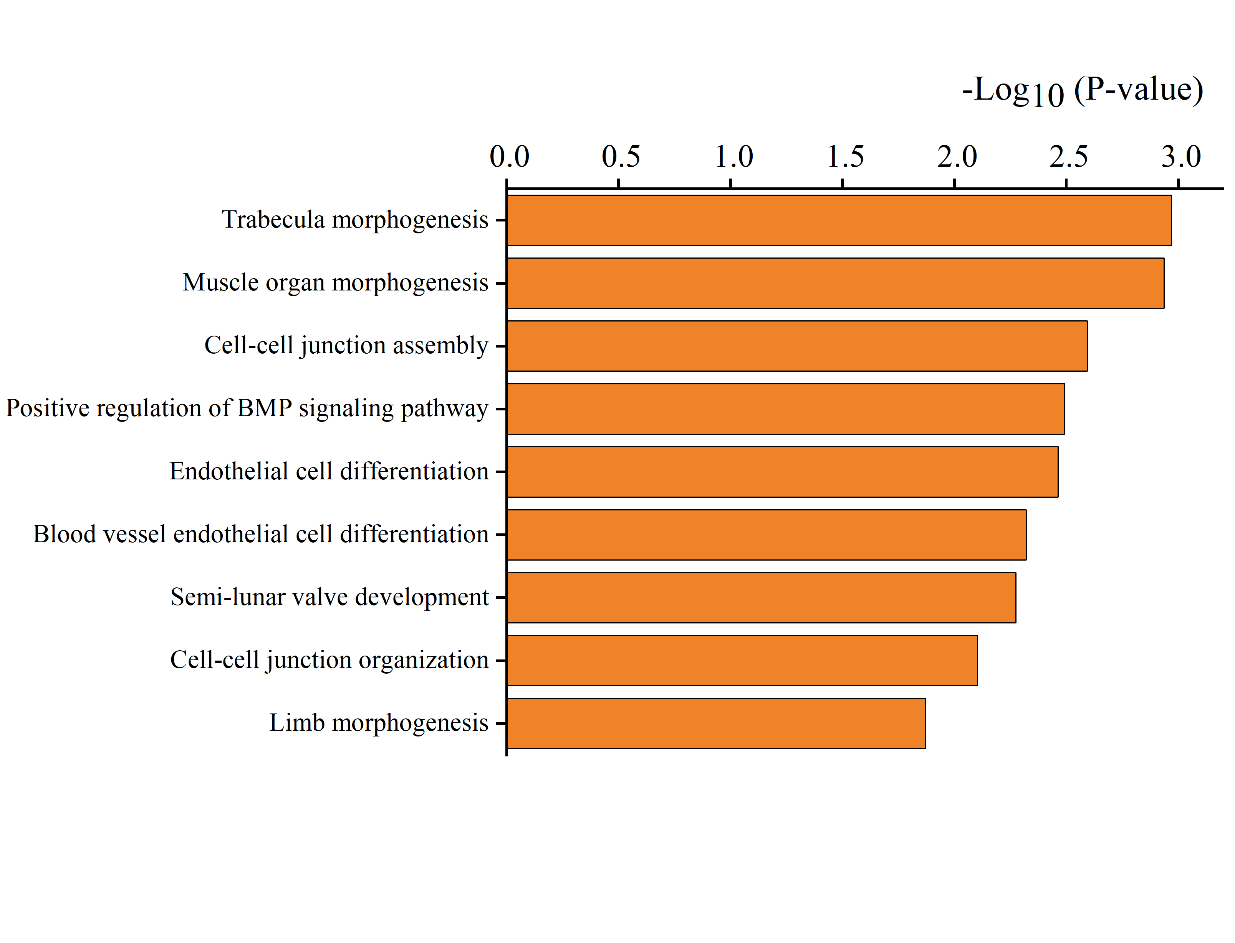

**Figure S12.** **Gene ontology enrichment related to biological process on the upregulated proteins of osteochondral interface compared with SB tissues.**

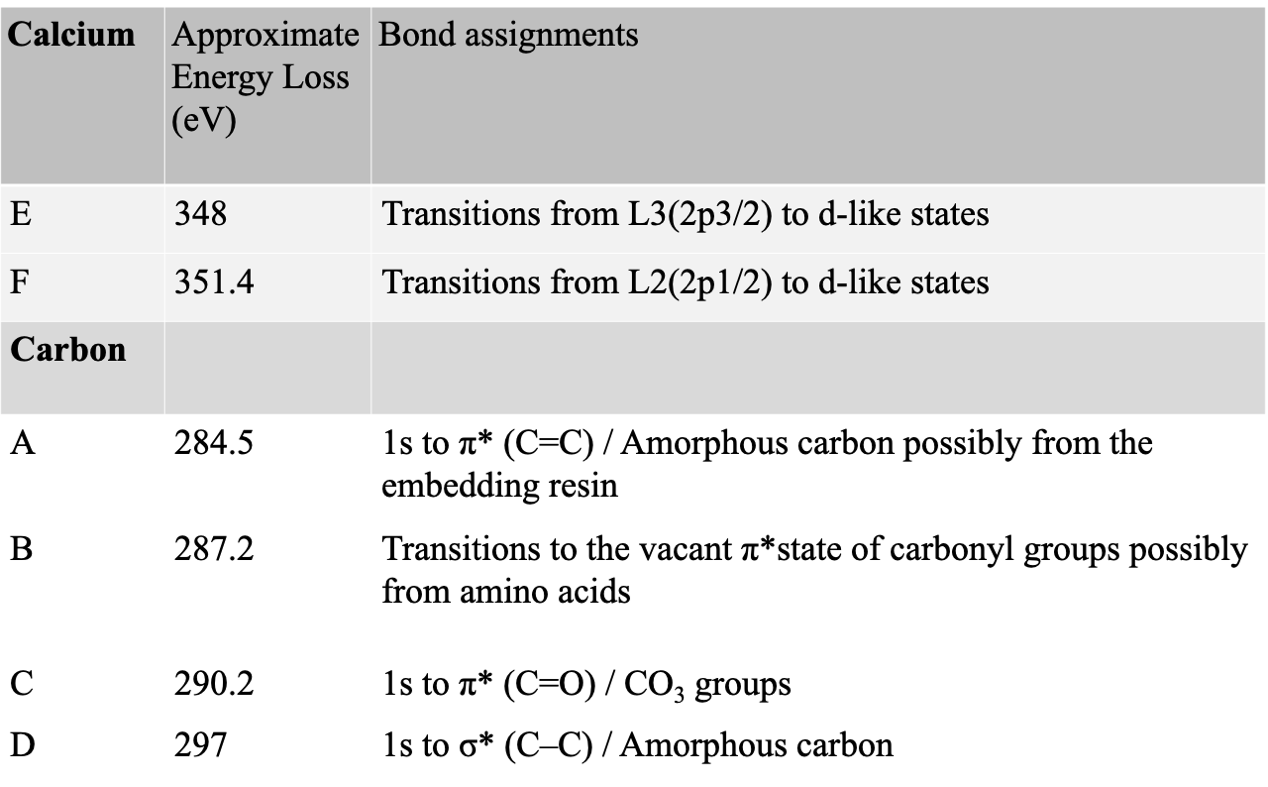

**Table S1.** **Electron energy loss spectroscopy (EELS)**. Approximate transitions in calcium L_2,3_ edge and carbon K edge fine structure and assignments of peaks.
